# scChat: A Large Language Model-Powered Co-Pilot for Contextualized Single-Cell RNA Sequencing Analysis

**DOI:** 10.1101/2024.10.01.616063

**Authors:** Hsuan-Han Chiu, Ashley Varghese, Kunming Shao, Yen-Chun Lu, Rahul Nahar, Hao Chen, Qing Deng, Xiaoping Bao, Can Li

**Author notes:** Corresponding authors (X. B.,; C.L.,). These authors contributed equally (Hsuan-Han Chiu, Ashley Varghese, Kunming Shao).

## Abstract

Single-cell RNA sequencing (scRNA-seq) has transformed biomedical research by enabling transcriptomic analysis at single-cell resolution. Yet, existing computational approaches remain primarily data-driven and lack the ability to integrate research context, limiting their interpretability and impact on hypothesis generation or experimental planning. We present scChat, a large language model (LLM)–powered co-pilot for contextualized scRNA-seq analysis. Unlike conventional pipelines restricted to tasks such as cell type annotation or enrichment analysis, scChat has an interactive, reasoning-based framework. It combines quantitative algorithms with retrieval-augmented generation and a multi-agent architecture to support hypothesis validation, mechanistic interpretation, and next-step experimental design. Through showcase and benchmarking studies, we demonstrate that scChat not only achieves high accuracy in cell type annotation but also provides biologically grounded explanations and contextual insights.

## Introduction

Single-cell RNA sequencing (scRNA-seq) has revolutionized biomedical research by providing a detailed and high-resolution view of the transcriptomic landscape within individual cells^1^. This technology enables researchers to dissect tissue heterogeneity, uncover distinct cell populations, and trace developmental or disease trajectories with unparalleled cellular resolution. Various machine learning algorithms have been developed to analyze scRNA-seq data, including unsupervised clustering methods implemented in software like Seurat^2^ and Scanpy^3^, as well as supervised^4–10^ and self-supervised learning algorithms^11–14^. Deep learning methods, particularly those employing transformer-based architectures like scBERT^11^, scFoundation^14^, and scGPT^12^have achieved impressive accuracy in tasks such as cell type annotation. However, a significant limitation of these deep learning approaches is their lack of interpretability, which hinders their usability for biomedical researchers without a computer science background. This barrier restricts interactive tasks such as obtaining biological insights from cell classifications and analyzing cell populations. Furthermore, the purely data-driven nature of ML analysis in scRNA-seq often fails to “contextualize” the specific biomedical problems under investigation, thereby limiting its ability to directly inform experimental design and hypothesis generation.

Large Language Models (LLMs), such as OpenAI’s GPT-4^15^ (the LLM powering ChatGPT) and Meta’s Llama models^16^, are trained on extensive datasets covering a broad spectrum of human knowledge. These pretrained LLMs can execute tasks, such as code generation, exam-taking, and essay writing at an unprecedented level. To harness the potential of LLMs, various “copilot” tools (AI assistants) have been developed based on these models. For instance, LLaVA-Med^17^, BiomedGPT^18^, Med-Gemini^19^ are chatbots to help doctors analyze medical images and generate diagnosis reports. Aside from commercial use, this idea has been adopted as a “co-scientist” in the scientific field, aiming to facilitate progress in research. Coscientist^20^, an AI copilot for chemical research powered by the OpenAI GPT-4 model, can autonomously design, plan and perform complex experiments. Similarly, the AI-Coscientist, presented by Gottweis et al. ^21^, shortened the timeline for developing the capsid-forming phage-inducible chromosomal islands hypothesis from twelve years to just two days. The GPT-4 model was also empirically tested to annotate cell types and was shown to perform close to human experts^22^. Together, these examples underscore the potential of AI-driven systems to accelerate scientific discovery significantly. However, a significant barrier to application in biomedical science and scientific research is the phenomenon of “hallucination”^23^, where an LLM might present information with unwarranted confidence, even when uncertain. Moreover, pretrained models like GPT-4 excel at qualitative tasks such as explaining technical concepts but struggle with quantitative tasks^24^.

To address these challenges, we introduce scChat, a platform designed for scRNA-seq analysis that emphasizes contextualized reasoning through a multi-agent architecture and extensive retrieval-augmented generation (RAG)^25^ techniques **(Fig. 1)**. Foundation models such as scGPT focus on learning gene embeddings from large-scale datasets to enhance downstream tasks like clustering and annotation, but typically require substantial computational resources and coding expertise. In contrast, scChat is positioned within the emerging class of conversational and workflow-oriented agents for omics analysis. Rather than training new foundation models, scChat prioritizes accessibility, interpretability, and automation of pretrained LLMs and established scRNA-seq workflows. Serving as a generalizable interactive co-pilot for scientists, scChat enables users to perform scRNA-seq data processing, cell type annotation, gene enrichment analysis, and statistical sample comparison through natural language interaction via a graphical user interface (GUI). Beyond these core functions, scChat leverages LLM-driven reasoning and structured function calls to address a wide range of analytical questions that arise during single-cell data analysis, thereby generalizing and unifying existing scRNA-seq analysis pipelines. Importantly, scChat achieves high accuracy in cell type annotation while explicitly explaining the role of marker genes, evaluating differential gene expression and pathway enrichment across conditions, and synthesizing diverse biological evidence to generate mechanistic insights. In this way, scChat extends traditional analysis pipelines into an interactive assistant that supports hypothesis verification, reduces hallucinations through grounded retrieval, and preserves the accuracy of quantitative analyses. Because its annotation and reasoning capabilities are grounded in retrieval-augmented generation, scChat can continuously improve as its underlying knowledge base evolves.

**Figure 1.**
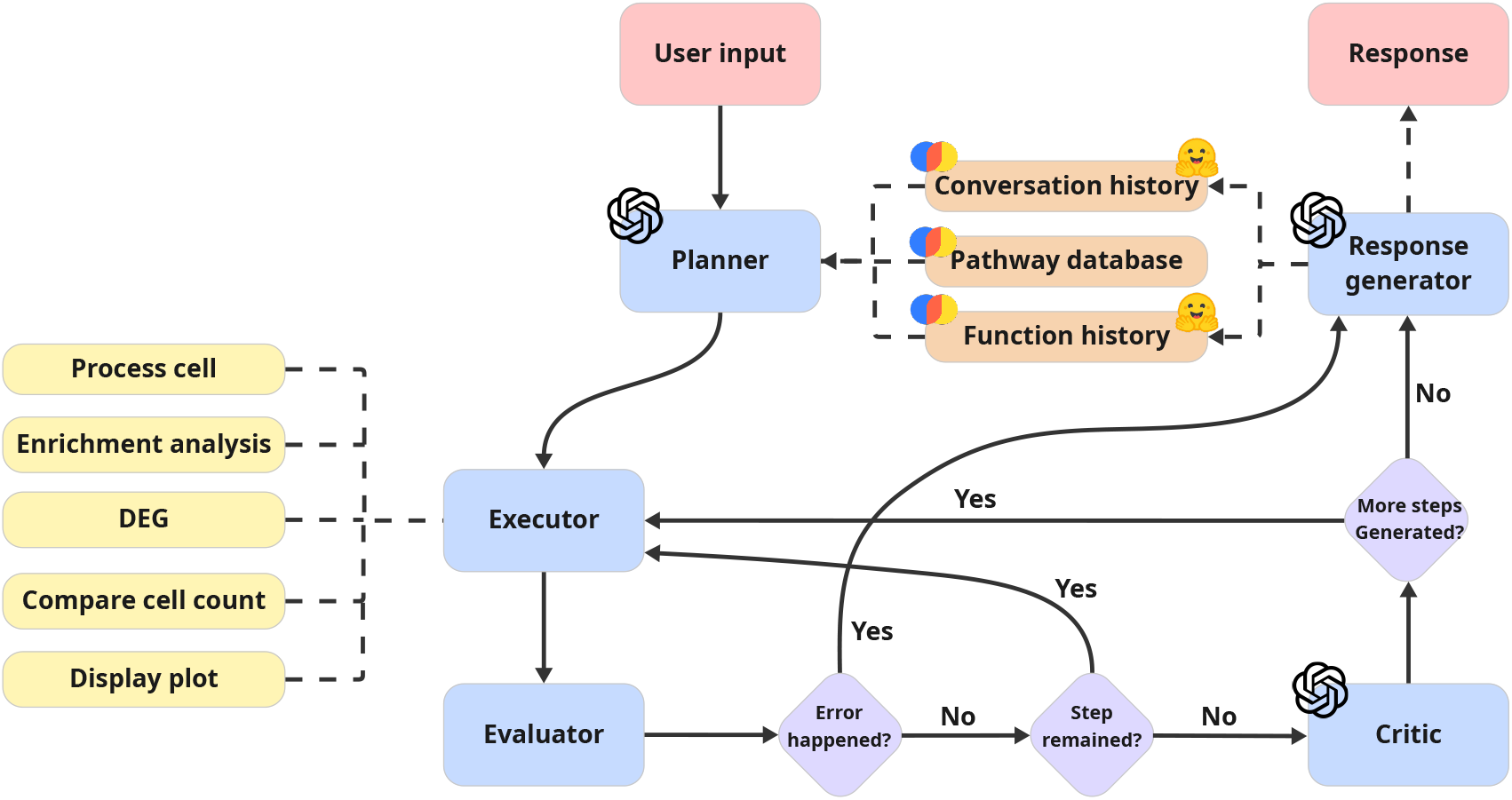
An overview of scChat multi-agent system framework.

### scChat Architecture Overview

**Fig. 1** provides an overview of scChat. As a multi-agent system (MAS), scChat includes five main agents: planner, executor, evaluator, critic, and response generator. The agents in scChat can use predefined functions as tools through a function calling process. When given raw scRNA-seq data and a user query via a GUI, the planner first reviews the conversation history and function history to retrieve relevant results, then creates a plan outlining all necessary analyses as steps involving function calls. Next, the executor collaborates with the evaluator to perform the functions and verify the availability of the results. All scRNA analyses in this system are performed using functions from the executor. The cell type annotation function uses cell type RAG to identify cell type level by level. The subsequent analysis, including differential gene expression (DEG) analysis, enrichment analysis, and comparison of cell type populations, is then conducted based on the identified cell types. Finally, scChat displays the analysis results via various kinds of plots. During execution, the evaluator plays a crucial role in error management by removing steps involving invalid cell types that are absent in the dataset from the plan and capturing all error messages, which are stored in the function history. The critic determines whether follow-up analyses are necessary for specific cell types in the user query. Lastly, the response generator assembles analysis results and responds to the user query. It stores responses and function execution histories in two vector databases, the conversation history and function history, which assist the planner in generating follow-up plans based on past analysis and conversation data.

## Results

### Showcase study

The showcase study demonstrates the interaction between users and scChat using raw single-cell sequencing data. To showcase the capabilities of our approach, we deliberately selected a highly complex human clinical trial dataset, as clinical trials inherently involve far greater biological and analytical complexity than *in vitro* systems or murine *in vivo* models, particularly due to inter-patient heterogeneity in treatment responses and clinical outcomes. Accordingly, we chose the study by Bagley et al. ^26^ as our showcase example. This study investigated whether adding pembrolizumab, a PD-1 inhibitor, could boost EGFRvIII CAR T therapy in glioblastoma. Specifically, based on previous findings that CAR T treatment may lead to PD-1 upregulation in the tumor microenvironment. As a result, the researchers investigated whether blocking the upregulation of PD-1 could prevent T cell suppression and enhance the effectiveness of CAR T cells. The testing samples were collected from seven adults with newly diagnosed EGFRvIII glioblastoma, including both pre- and post-treatment conditions. This testing highlights scChat’s potential to support hypothesis refinement and contextual interpretation of scientific findings.

In the original clinical trial, no significant therapeutic benefit was observed at the cohort level, with a median progression-free survival of only 5.2 months and an overall survival of 11.8 months. CAR-T cells exhibited limited expansion and persistence in peripheral blood, and CAR-related fragments were detected only transiently in the tumors of a few patients. We sought to evaluate whether scChat could recapitulate key findings directly from the raw single-cell data, thereby validating its utility as a research co-pilot. As this dataset derives from brain tumors, one of the intended goals of CAR-T infusion was to remodel the immunosuppressive tumor microenvironment (TME). Moreover, since samples underwent CD45 magnetic enrichment before scRNA-seq, the majority of cells were expected to be immune cells of peripheral origin. We therefore first instructed scChat to perform fine-grained clustering and annotation of immune cells.

As shown in **Fig. 2a**, scChat offers an interactive graphical interface that enables users to upload data. Following preprocessing, users can issue queries. In **Fig. 2b**, scChat annotated immune cell populations in response to prompts and visualized them with UMAP, reconstructing an immune landscape of the TME. Out of 11,424 cells from six samples, including pre- and post-treatment of P1, P6, and P7, the system identified 26 clusters, consisting of both immune and a small number of non-immune cells. The TME was dominated by macrophages and T cells, with relatively few B cells. Notably, consistent with the original study, scChat also identified brain-resident immune cells, such as microglia, which are characterized by low CD45 expression. Detailed annotations are provided in **Fig. 2c**.

**Figure 2.**
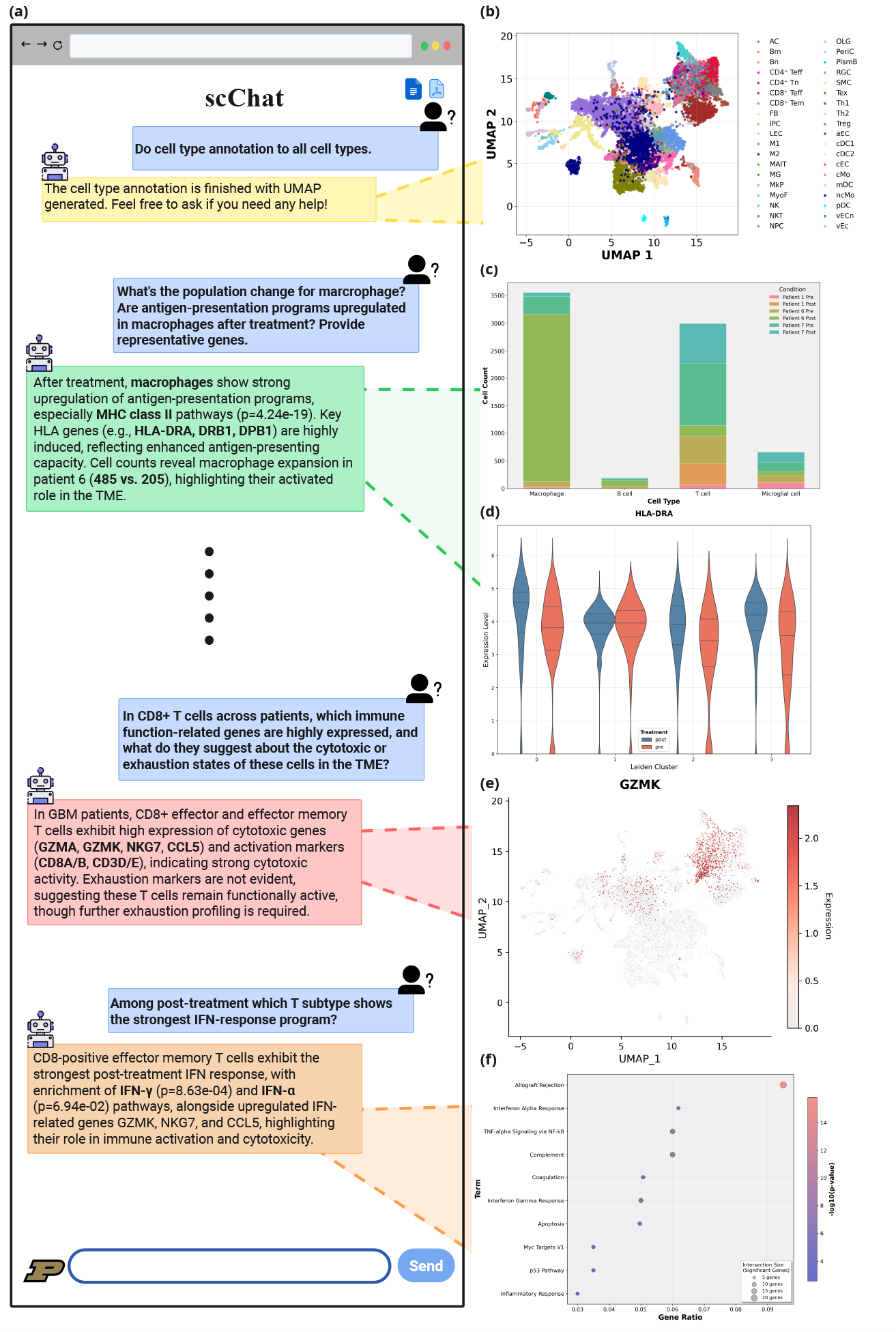
scChat-assisted analysis of immune composition in the tumor microenvironment with therapy. (a) Representative snapshot of the scChat interactive conversation window. (b) UMAP visualizes the overall cellular landscape of the tumor microenvironment (TME) from three paired patients (pre- and post-CAR T-cell infusion from patients 1, 6, and 7; six samples in total). (c) Relative abundance of each cluster in each patient sample. (d) Violin plots showing HLA-DRA expression across clusters in Pre versus Post samples. (e) Feature plot of GZMK expression. (f) Gene set enrichment analysis (GSEA) for the cluster of interferon-stimulated T cells, with multiple testing correction using the Benjamini–Hochberg procedure.

We next asked scChat to compare pre- and post-treatment samples. The system not only reported cell-type proportions per patient, but also visualized compositional changes. Patient P1 displayed an increase in T cell proportion following therapy, whereas patients P6 and P7 exhibited a marked expansion of macrophages. Interestingly, P1 had a superior clinical outcome compared with P6 and P7, suggesting that tumor-associated macrophage (TAM) accumulation may correlate with poor prognosis and play a central role in the TME.

To investigate how innate and adaptive immune compartments were qualitatively affected by treatment, we instructed scChat to separately subcluster T cells and macrophages. The analysis identified 10 T cell subclusters and three macrophage-like subclusters, as shown in **Fig. S1a**. Although the annotations did not exactly match those in the original study, which defined populations such as TREM2^hi microglia using high-expressing marker genes, we observed that these represent user-defined subtypes not directly recognized by scChat. To address this, we further queried scChat regarding the dynamics of microglia and macrophages before and after therapy. Through differential gene expression and population analysis, the system detected behaviors in these populations consistent with those reported for TREM2-high microglia and monocyte-derived macrophages, accurately reflecting patient-specific changes. As scChat concluded, there was a decrease in TREM2-high microglia and an increase in monocyte-derived macrophages in P1 only.

Notably, Alanio et al. previously reported that the presence of IFN-stimulated T cells at relapse serves as a significant prognostic biomarker in GBM^27^. Accordingly, we focused on IFN-related immune cells in our dataset to validate this finding. As a co-researcher, scChat not only analyzes data and provides feedback but also engages with the user to suggest research directions and guide next steps. Given scChat’s identification of macrophages as antigen-presenting cells in the samples, as displayed in **Fig. 2d**, we explored their potential interactions with T cells after treatment. The observed upregulation of MHC-II in macrophages and increased interferon-stimulated gene (ISG) expression in T cells, along with scChat’s inference shown in **Fig. 2e**, suggest a synergistic interaction between these two cell populations, where macrophage antigen presentation may enhance interferon-driven T cell activity^28^. Macrophages can activate helper T cells through MHC-II–mediated antigen presentation, and in turn, these T cells secrete cytokines that enhance macrophage function and further promote immune activation. This is supported by gene expression evidence, such as the upregulation of CCL3 in macrophages, which can recruit and activate T cells. Guided by this framework, we next queried scChat about the expression of antigen-presentation and interferon-stimulated gene (ISG) programs in macrophages before and after therapy. Since myeloid cells such as TAMs secrete chemokines and suppressive cytokines that directly influence T cell activation and exhaustion, and given that scChat had already indicated remodeling of the myeloid compartment, as shown in extended data, in the TME, research has previously reported that although T cells are recruited into the tumor, without sustained antigen stimulation, they may eventually progress toward exhaustion.

We hypothesized that similar changes would also occur in T cells. We therefore asked scChat to analyze isolated T cell subsets, including CD8^+^, CD4^+^, and Tregs. scChat successfully identified CD4^+^ effector T cells, CD8^+^ effector T cells, NK cells, T helper cells, and additionally annotated a subset of exhausted T cells, which is consistent with the expected coexistence of activated and exhausted T cell states in the TME. When requested, scChat generated violin and feature plots of canonical exhaustion markers, confirming increased expression after treatment. Based on the macrophage antigen-presenting and ISG signatures described above, scChat inferred that T cells would also show stronger interferon-stimulated programs post-therapy. This was validated by analysis showing that CD8^+^ effector T cells exhibited the strongest IFN-response signature after treatment, supported by GSEA evidence of significant enrichment, which is included in **Fig. 2f**.

Finally, scChat integrated the observed changes in the myeloid and T cell compartments into a coherent mechanistic model; both macrophages and T cells within the same TME showed enrichment of interferon pathways and upregulation of ISGs, indicating potential coordinated activation. In this model, T cell– derived interferons activate macrophages, polarizing them to the antigen-presenting states. In contrast, macrophages support T cell function by presenting antigens and releasing inflammatory cytokines, thereby creating a positive feedback loop that enhances T cell activity. This dynamic not only illustrates the immune circuitry of the TME but also offers a mechanistic explanation for the synergistic antitumor effects seen with CAR-T and PD-1 blockade. Additional details of the scChat conversation underlying this EGFRvIII-directed CAR-T combination therapy study are provided in **Fig. S1**-**S2** and the summarized Q&A can be found in **Table S1**. These findings align with those from the original study, highlighting that scChat can serve as an effective co-pilot for biomedical research, helping with data processing, visualization, logical reasoning, and hypothesis generation.

### Cell type annotation accuracy analysis

As another case study, we evaluated the accuracy of cell type annotation using eight datasets from Xu et al.’s ^29^ study. Specifically, the organ and tissue atlases include cells from the lymph node, pancreas, kidney, blood, bone marrow, spleen, and liver. As a key function of scRNA-seq analysis, this assessment aims to measure the performance of cell type annotation from scChat; however, the cell type names may vary across databases, making scoring difficult and leading to a semantic matching issue. To avoid the problem of cell type name mismatches, we use the cell ontology from the CellxGene platform^30^ as the ground truth cell type lineage. By mapping cell type names predicted by different models to this unified naming system, we then evaluate performance using the hierarchical F1 score (hF1). Briefly, hF1 is defined in terms of precision and recall, as described in the Supplementary Information **equations (1)-(8)**. With respect to the cell type ontology, precision and recall assess whether an annotated cell type is overly specific or overly broad, respectively. In particular, predictions that are too broad lead to reduced recall, whereas overly specific predictions result in lower precision. The hF1 metric integrates both precision and recall providing a balanced evaluation of annotation performance. The explanation of the annotation pipeline and the hF1 calculation is detailed in the method section. Six models, including SingleR ^31^, GPT-4o, CellTypist^32^, Azimuth^33^, scGPT^11^, and scChat, are tested with the same datasets. Notably, the GPT-4o here applies the same data preprocessing strategy as scChat but without the support of cell type RAG. Each model takes the h5ad file as input and outputs predicted cell types for every barcode. While the hierarchical F1 score provides a rigorous quantitative benchmark, we further visualize predictions using UMAP to reveal how well each model preserves biological structure and to make differences more interpretable through a spatial view of annotation. **Fig. 3** displays the UMAPs from all the tested models with blood tissue. It is evident that the visualizations from scChat and scGPT exhibit finer granularity compared to those from SingleR and GPT-4o. This supports the finding that both scChat and scGPT achieve higher performance in hF1, as discussed in the following analysis. Notably, the rest of the comparison results are shown in **Fig. S3-9**.

**Figure 3.**
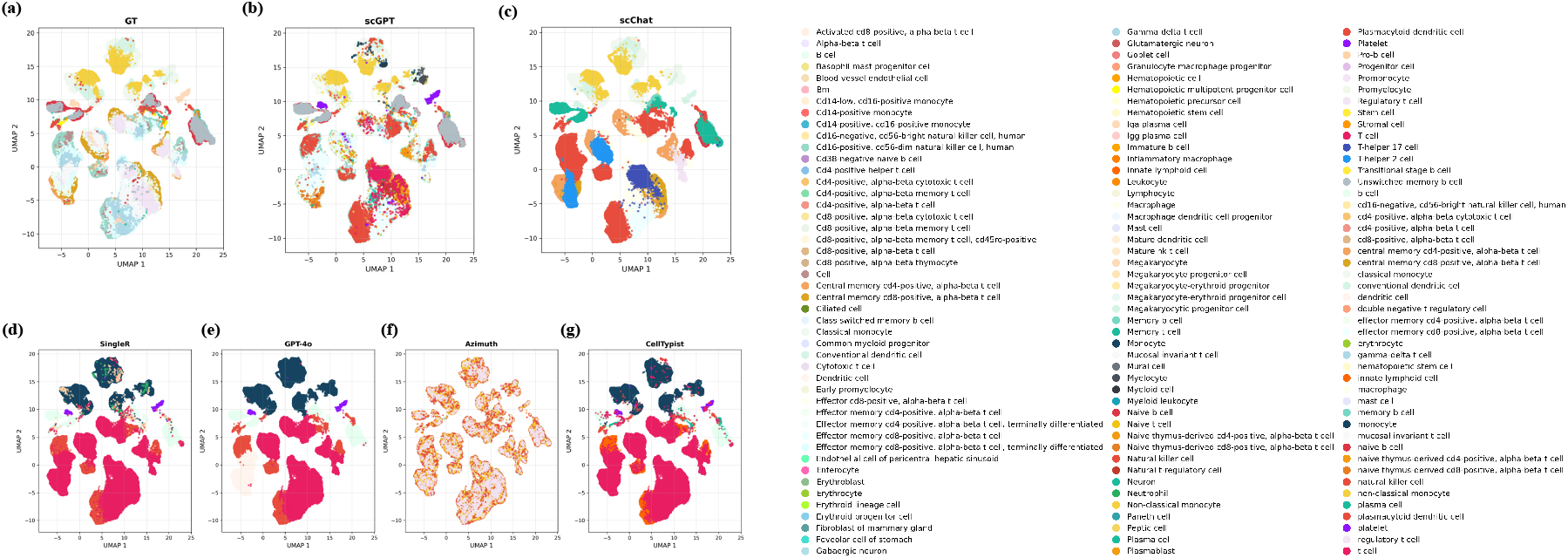
UMAP for cell type annotation results. (a) Ground truth (GT) from the original dataset. (b) scGPT. (c) scChat. (d) SingleR. (e) GPT-4o. (f) Azimuth. (g)CellTypeist.

**Fig. 4a** shows the hF1, precision, and recall for all models. In our study, scChat demonstrates outstanding performance in cell type annotation across all tested datasets. It consistently outperforms GPT-4o and SingleR, CellTypist, Azimuth, while achieving results comparable to scGPT, the state-of-the-art pre-trained model. Specifically, scChat achieves a 0.886 hF1 score in the intestine dataset, surpassing the 0.843 score of scGPT. The hierarchical precision and recall in **Fig. 4** reveal a trend that most models perform better in precision than recall across all datasets. This suggests that models, especially GPT-4o, SingleR, and CellTypist, tend to make fewer specific annotations. When unsure about the exact subtype, models like GPT-4o, SingleR, and CellTypist often default to broader cell categories, such as general cells, T cells, or B cells. The tendency toward broader cell type annotation can be traced to differences in how models handle the marker gene. Azimuth exhibits a more balanced performance between precision and accuracy and demonstrates superior performance on the kidney dataset. These observations can be attributed to two main reasons. First, Azimuth’s predefined models contain fewer hierarchical cell type levels, typically no more than three. In addition, the cell types defined at these levels are more closely aligned with the ground truth annotations in the testing atlases. As a result, Azimuth is able to achieve a favorable balance between precision and accuracy. However, in real-world scenarios, this design may limit its ability to predict intermediate or fine-grained cell types. Second, Azimuth currently does not provide pretrained models for spleen, intestine, or lymph node tissues, and only limited information is available for kidney. Consequently, for these tissues and organs, we applied a 5-fold cross-validation strategy, in which each dataset was split into 80% training data and 20% testing data, repeated over five rounds to generate predictions across the full dataset. This evaluation setting likely leads to an apparent overfitting behavior in the kidney dataset, contributing to Azimuth’s inflated performance in this case. scGPT is trained to focus on the full range of expressive genes, rather than mainly relying on the top-expressed housekeeping genes. In contrast, scChat explicitly uses RAG to exclude housekeeping genes during the annotation process. These mechanisms enable both scGPT and scChat to produce more detailed predictions, whereas SingleR and GPT-4o often focus on housekeeping genes, resulting in less granular predictions. While this approach increases precision by avoiding misclassification, it does so at the cost of recall and reduces the biological resolution of the annotation. In comparison, scGPT and scChat maintain a more balanced precision and recall profile. For example, the results from the bone marrow dataset show that scChat has a difference of about 0.002 between precision and recall, which is much better than the differences of 0.118 and 0.074 from singleR and GPT-4o. We attribute scChat’s performance here to the cell type RAG. The complex cell lineages enable scChat to filter out less critical housekeeping genes and identify specific cell types more effectively. In conclusion, this study demonstrates that augmenting a general LLM with a domain-specific RAG can match or even surpass the performance of a fine-tuned model. In other words, delivering target information through RAG is a powerful, training-free strategy to close the performance gap compared to a well-trained foundation model.

**Figure 4.**
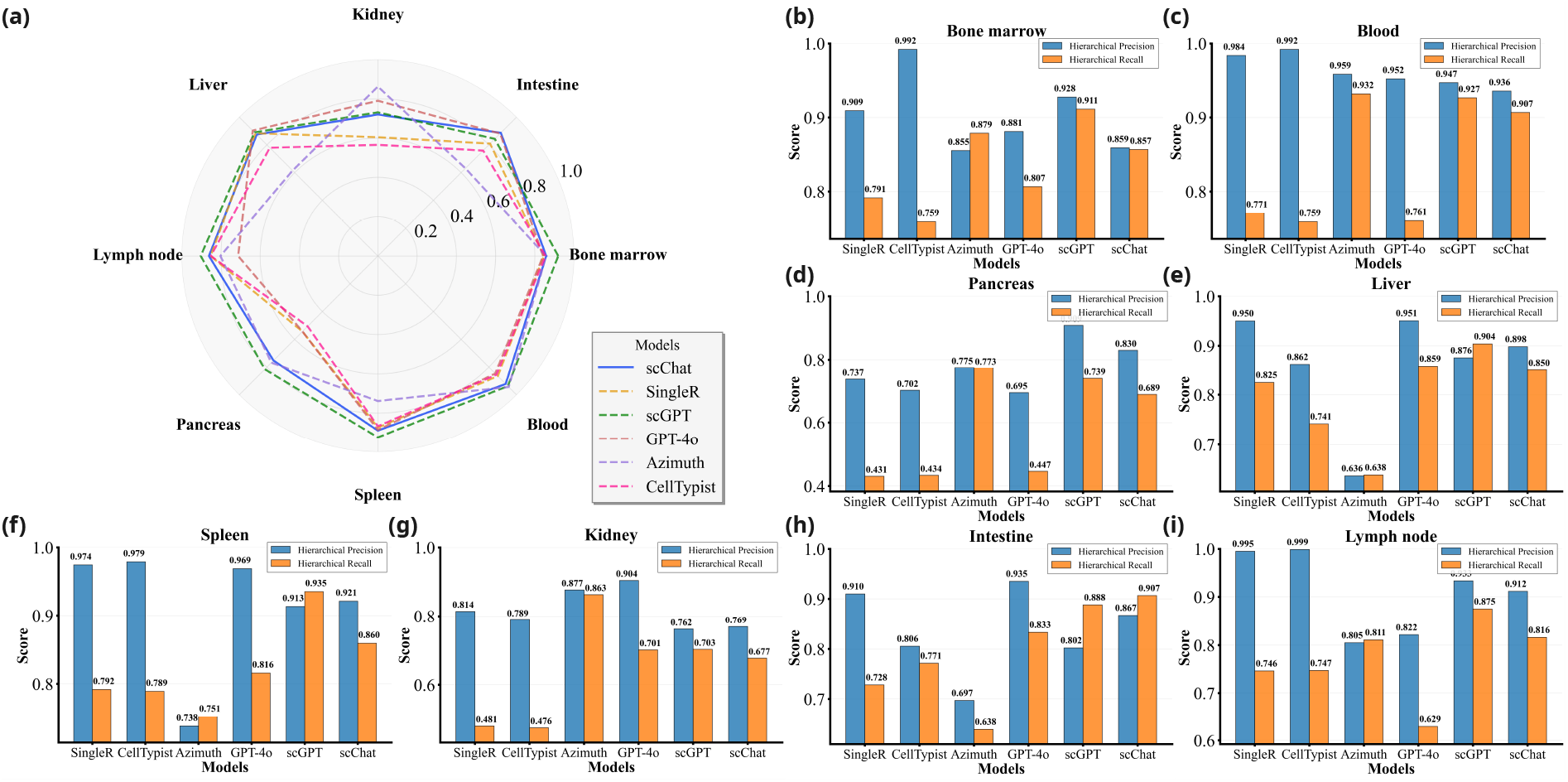
Cell type annotation accuracy analysis, including hierarchical F1-score in the radar plot and hierarchical precision and recall in the line plots.

### Question benchmarking

To systematically evaluate the performance of scChat, we established a benchmarking question set with four categories. The question set is generated from eight journal articles^26,34–40^. These datasets were selected to encompass a broad range of research contexts and organ systems, allowing us to thoroughly evaluate scChat in various biological settings. We classify the queries to scChat into four categories: cell type identification, differential gene expression analysis, gene enrichment analysis, and complex queries. Additionally, we have 43 questions for cell type identification, 26 questions from differential gene expression analysis, 15 questions for enrichment analysis, and 27 questions for complex queries. After testing, GPT-4o evaluates scChat’s responses using a strict evaluation rubric and generates scores along with corresponding rationales. Bio-experts then review the scoring rationales to verify the appropriateness of the assigned scores. The full question-based benchmarking pipeline is illustrated in **Fig. 5. Table 1** presents the objectives and examples of each category we have created. The usage of LLM for question generation and evaluation is outlined in the method section.

**Figure 5.**
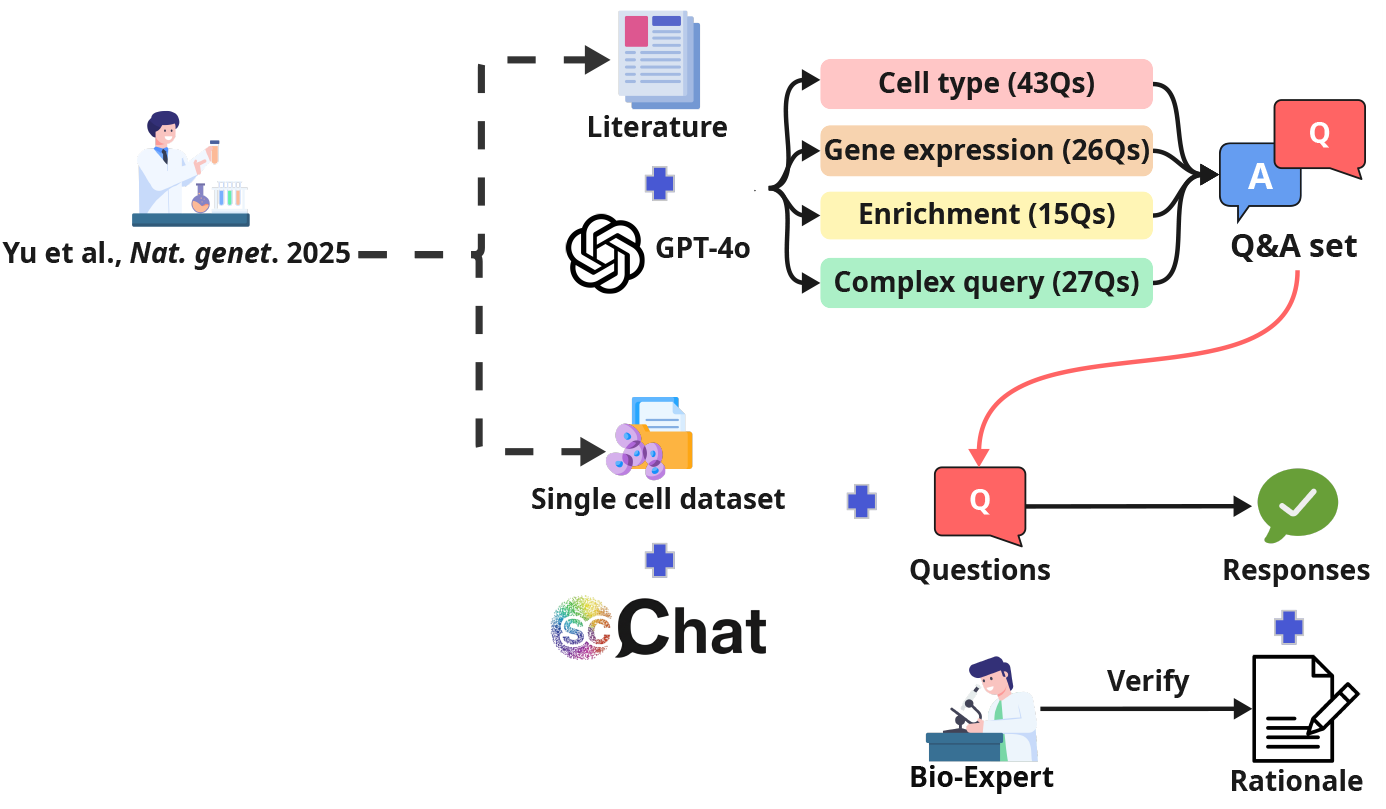
Demonstration of question-based benchmarking pipeline.

**Table 1.** Categories of the benchmarking question.

|  | Objective | Example of query |
| --- | --- | --- |
| Cell type identification | Test the functionality to find target cell type. | How can we differentiate between hematopoietic stem cell and T cell using their canonical markers? |
| Differential gene expression | Measure the scChat with its capability to observe the changing in specified gene. | Regarding hematopoietic stem cells, what changes in CD34 expression were observed when comparing young healthy individuals to aged healthy ones? |
| Enrichment analysis | Evaluate the scChat's biological understanding to choose correct database for enrichment analysis. | Can we confirm if the Immune Response is enriched in T cells from the young healthy? |
| Complex query | Assess the chaining of multiple functions to answer the user query. | Based on the successful identification of T cells, the enrichment of the IFN-stimulated gene response in influenza, and the expression changes in COVID, TNF, IL, synthesize a comprehensive mechanistic explanation for how these elements interact to drive the observed therapeutic response across pre-treatment, post-treatment. |

For the cell type identification question, scChat recognizes the targeted cell type mentioned in the query and clarifies the finding with the marker gene and rationale to answer the question. Using the question in **Table 1** as an example. To answer this correctly, scChat first acknowledges that the question asks about hematopoietic stem cells and T cells. It further notes that in the dataset, hematopoietic stem cells are annotated with upregulated PROM1, MPL, and FLT3. T cells, on the other hand, are annotated with the CD3 complex, CD4, CD8A/B, TRAC, and IL7R. By comparing the marker genes used for each cell type, scChat achieves a perfect 5/5 score for this question and an average score of 4.72/5 for all the cell type identification questions, as shown in **Fig. 6**.

**Figure 6.**
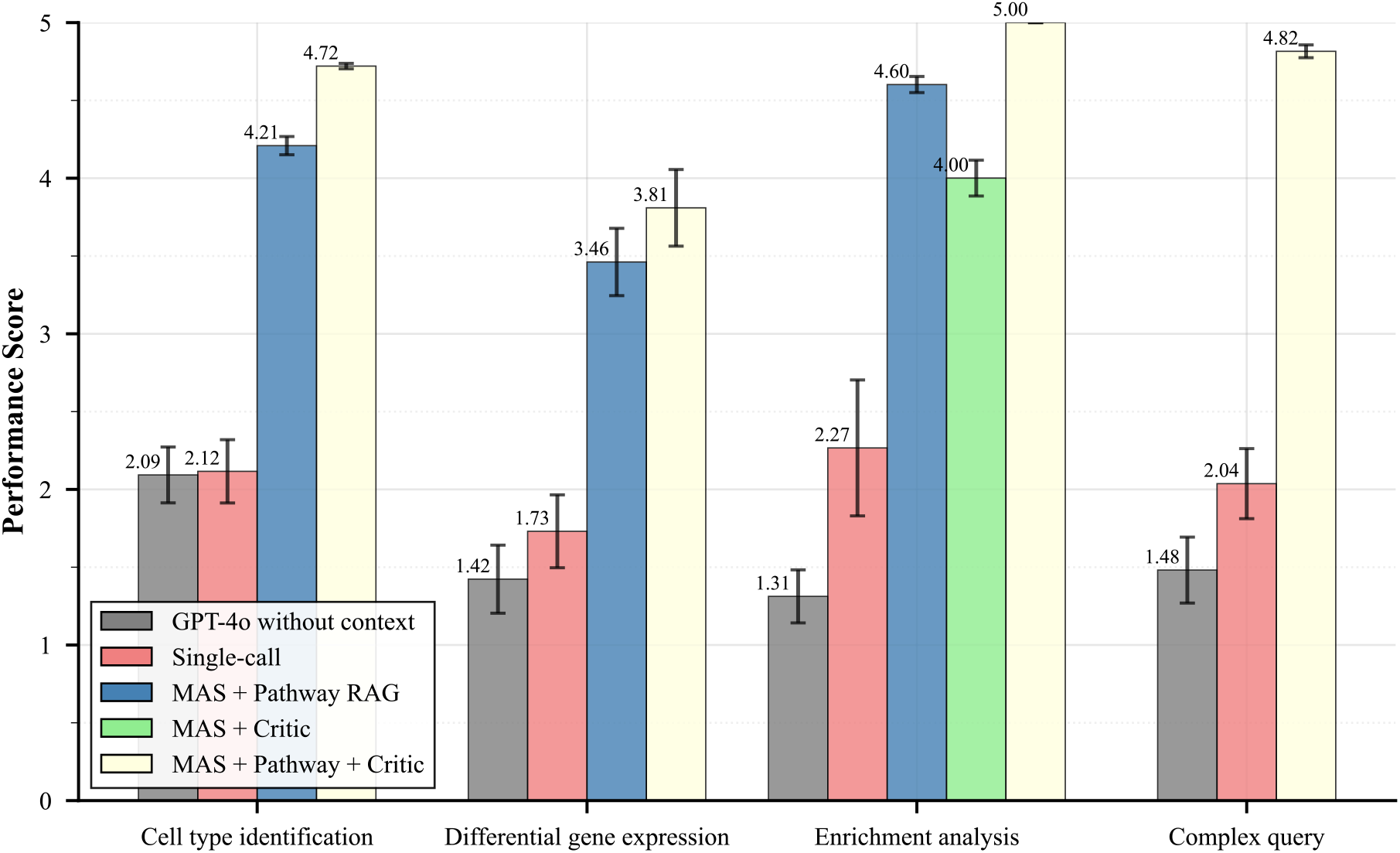
Performance of each system. GPT-4o is used to evaluate the responses shown in this plot, following clear evaluation instructions and a scoring rubric. Based on the instructions and rubric, GPT-4o assigns a score and provides a corresponding rationale for each response. A domain expert then reviews the rationales to verify that the assigned scores are reasonable. With the assistance of the pathway RAG and revision loop, the final version of scChat performs the best across all categories of benchmarking questions. During failure mode testing, we examine the rationale for not achieving a full score in each category. We also record the performance from systems with functionality removed for an ablation study.

The differential gene expression question prompts scChat to conduct a follow-up analysis and confirm the change in gene expression or the observation of a specific upregulated gene. As to the question in **Table 1**, scChat performs DEG on hematopoietic stem cells and identifies the difference in CD34 expression between young and aged samples. ScChat finds that CD34 remains highly expressed in both conditions, which is consistent with the findings from ground truth. Therefore, ScChat receives a 5/5 score for this question. In this type of question, scChat gets an average score of 3.81/5.

Similar to the question about differential gene expression, the enrichment analysis question requires scChat to perform downstream analysis to provide a response. To answer the question from **Table 1**, scChat conducts enrichment analysis on T cells and retrieves pathways such as T cell activation, lymphocyte activation, and immune system processes from GO, and T cell receptor signaling pathway and cytokine-cytokine receptor interaction from KEGG, demonstrating the robust immune response that T cells are experiencing. This question ends up achieving a 5/5 score, and scChat receives an average score of 5/5 for this type of questions.

Lastly, scChat performs a chain of functions to gather all necessary information to answer the complex queries. To address the question in **Table 1**, scChat first conducts an enrichment analysis on T cells. The results indicate that TNF-α signaling via the NF-κB pathway is significantly enriched, suggesting a key role in amplifying pro-inflammatory cytokine production. While the differential gene expression analysis highlights strong upregulation of IL7R and IL32 in COVID-19-affected patients, this suggests survival and activation of cells. It concludes that targeting TNF and IL-1β in COVID-19 may reduce hyperinflammation, while modulating IFN-I responses, enhancing their early antiviral capacity, and restraining their late pro-inflammatory effects, could improve disease outcomes. The distinct immune profiles between influenza and COVID-19 underscore the need for tailored therapies that balance viral defense with inflammation control. This response also scores a 5/5 for this question, and scChat receives an average score of 4.82/5 in all the complex queries.

### Failure mode testing

In failure mode testing, we review all imperfect responses from the evaluated results displayed in **Fig. 6** and categorize the reasons why scChat does not achieve a full score into three main groups.

First, inadequate analytical grounding includes responses with specific information, such as target cell types and genes that are not found. We can further attribute the causes of this type of failure to RAG mismatch and fundamental differences caused by the algorithm. In this system, the discovered cell types are strongly aligned with those in the cell type RAG. When the user queries a cell type that does not exist in the cell type RAG, it turns out to be a question beyond scChat’s capability. Additionally, various algorithms have aided researchers in scRNA-seq analysis, and different algorithms may yield distinct sets of expressed genes. This means scChat might identify the targeted cell type but fail to find the target marker genes. This group accounts for 36.4% of all incomplete responses, mostly within the differential gene expression category of benchmarking questions, resulting in a 3.81/5 scoring.

The second group involves deviations from canonical markers. This includes questions where the marker gene in the cell type RAG differs from what the user asks about or the different input system. For example, a question from the newborn lung cell dataset states that monocytes are annotated with CD11c, Tim3, and CD33. Although monocytes are correctly identified, none of these markers are used in the cell type RAG during cell type annotation. The question asks to annotate endothelial cells using CD31, which is supposed to be an antibody-based marker, whereas our RAG uses PECAM1, an RNA-based marker, instead. Questions in this group relate to cell type identification and make up 36.4% of all inadequate responses. According to the evaluation rubric, these responses are imperfect, but highlighting only the core finding results in a minor penalty and produces a score of 4.72/5.

The last issue concerns incomplete coverage of the question’s scope. scChat identifies each sample condition based on researcher-defined sample labels and additional descriptions. This allows users to add descriptions to each condition beyond the dataset’s basic labels, enriching scChat’s knowledge. However, some conditions mentioned in the questions are not included in the dataset. For example, when a question asks about both healthy and age-related macular degeneration patients, but the dataset only includes patients with age-related macular degeneration, this leads to insufficient information. All questions with such metadata mismatches fall into this category, which accounts for 27.2% of the flawed responses.

The first two issues could be addressed by transforming the cell type RAG system into an agentic RAG system. Instead of making the RAG a static database, allowing it to update information by searching for new cell types and marker genes can prevent the limitations of the current cell type RAG’s knowledge. However, the method to avoid hallucination of searched cell types and markers may become a core challenge for future research. The last issue, on the other hand, can be solved through a carefully user-defined description. As long as the user asks a question that aligns with the data and the sample condition descriptions, the issue can be resolved.

### Ablation study

To analyze the contribution of each module, we conducted an ablation study comparing multiple system configurations We note that this comparison isolates the effect of iterative planning and multi-step function execution, rather than serving as a general comparison between single-agent and multi-agent LLM systems. In total, five system frameworks were evaluated. The single-call system corresponds to a legacy scChat configuration in which all functionalities are invoked within a single model call. In contrast, the multi-agent system (MAS) equipped with both the pathway RAG and the revision loop represents the final version of scChat proposed in this study. Two additional ablated variants were also evaluated: an MAS with only the pathway RAG and an MAS with only the revision loop. The GPT-4o without context is the baseline in which queries were directly submitted to GPT-4o without additional analytical information. **Fig. 6** summarizes the performance of each framework. Within all the agents that can do function call, the single-call exhibits the lowest performance. This is primarily because many benchmarking questions require scChat to execute multiple analytical steps in a specific sequence. However, the single-call framework is limited to a single function invocation per query, which leads to incomplete analytical workflows, insufficient contextual information, and consequently lower benchmarking scores.

#### Question

The pathway analysis for the known ‘CD14+ monocytes (Monocytes)’ population from the COVID-19 patients displays what kind of key observation regarding the ‘IFN signaling’ signature, including its significance?

#### Ground truth

The pathway analysis for the ‘CD14+ monocytes’ population from COVID-19 patients shows that the ‘IFN signaling’ signature is activated in these cells, with heterogeneous intensity across different donors. This indicates variability in the interferon-stimulated gene (ISG) signatures among patients, which is a key observation in the study.

#### scChat

The pathway analysis for the ‘CD14+ monocytes’ population from COVID-19 patients reveals a significant enrichment in the ‘Interferon (IFN) signaling’ signature. Specifically, the Gene Ontology Biological Process (GO_BP) analysis shows a strong association with immune system processes, including the defense response (p=1.25e-19) and response to biotic stimulus (p=1.88e-17), which are indicative of heightened IFN signaling activity. This is further supported by the Reactome pathway analysis, where pathways related to the innate immune system and Toll-like receptor cascades are significantly enriched (p=2.67e-26 and p=3.61e-06, respectively). These findings suggest that CD14+ monocytes in COVID-19 patients are actively participating in an IFN-mediated immune response, which is a critical component of the body’s defense mechanism against viral infections like SARS-CoV-2. The significance of these pathways underscores the role of monocytes in mounting an antiviral response, potentially contributing to the inflammatory milieu observed in severe COVID-19 cases.

#### GPT-4o without context

In COVID-19 patients, pathway analysis of CD14+ monocytes often reveals an upregulated interferon (IFN) signaling signature. This is a significant observation because IFN signaling is a crucial component of the innate immune response to viral infections. In the context of COVID-19, this upregulation reflects the body’s attempt to combat the SARS-CoV-2 virus by activating antiviral defense mechanisms. The significance of this finding lies in the dual role of IFN signaling: while it helps in controlling viral replication, dysregulated or excessive IFN responses can contribute to inflammation and tissue damage, which are observed in severe COVID-19 cases. In some patients, an overactive IFN response can lead to a cytokine storm, exacerbating the condition. Conversely, a delayed or inadequate IFN response might lead to insufficient viral control, allowing the infection to progress. Therefore, understanding the IFN signaling signature in CD14+ monocytes can provide insights into the immune dysregulation observed in COVID-19 and help identify potential therapeutic targets to modulate the immune response for better clinical outcomes.

In **Fig. 7**, we compare the questions and corresponding responses from the ground truth, scChat, and GPT-4o without context. For the same question, GPT-4o without context is able to generate responses that are broadly consistent with immunology domain knowledge; however, its answers tend to remain general and primarily reiterate high-level immunological concepts. In contrast, scChat’s responses incorporate concrete supporting evidence derived from the execution of analytical functions, such as statistical testing and quantitative comparisons. This evidence-based reasoning explains why scChat achieves higher evaluation scores and underscores the importance of its ability to invoke and execute appropriate analytical functions to improve response quality.

**Figure 7.**
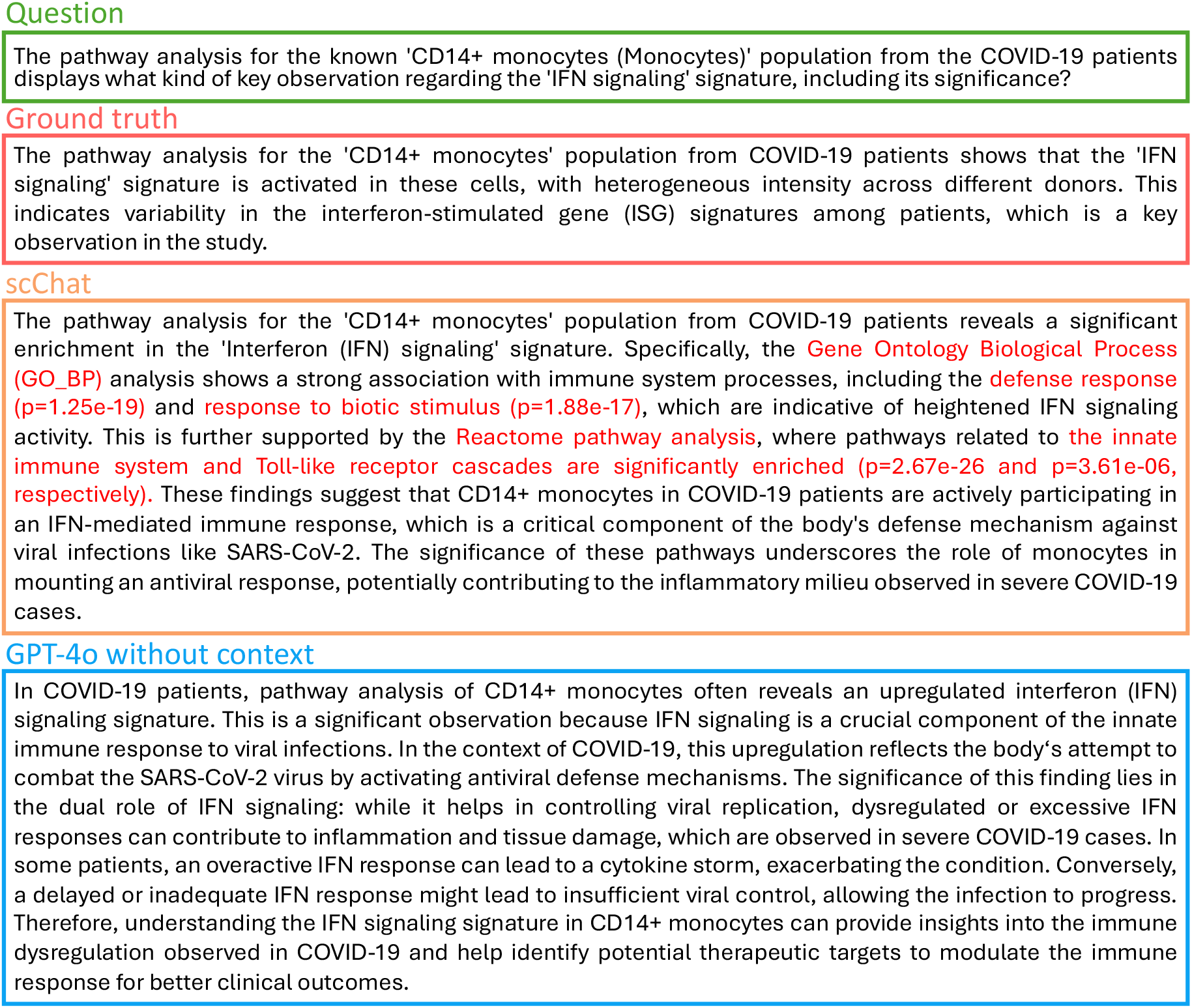
Comparison of results from the ground truth, scChat, and GPT-4o without contextual information. Keywords marked in red in scChat’s response indicate supporting information retrieved through function calls.

Comparing the MAS frameworks, systems with a revision loop can achieve better performance in cell type identification and differential gene expression categories. The revision loop appears in the critic, which categorizes questions into subordinate categories based on cell types mentioned in the query. Then, the critic creates a new plan that considers the subordinate category and function execution history. Unlike the original plan, which considers how to arrange the steps required to answer the question, the new plan focuses on steps for each cell type. This approach helps prevent missing analyses and enhances overall performance by approximately 0.5 average score in the first three categories.

The pathway RAG enhances scChat’s enrichment analysis by allowing the chatbot to dynamically select the most appropriate method, such as GO, KEGG, Reactome, or GSEA with its related gene set library, based on the user’s query. The analysis results, along with metadata, such as pathway names, term IDs, p-values, intersection sizes, and intersecting genes, are embedded into a vector database. This setup enables the semantic retrieval of analysis outcomes, rather than restricting results to default outputs, such as only the top-ranked terms. Consequently, pathway RAG reduces the risk of missing biologically relevant pathways and boosts the chatbot’s ability to match user search terms with suitable results. When combined with a revision loop, this approach consistently delivers optimal performance, achieving a perfect evaluation score of 5/5.

Overall, the revision loop enhances the thoroughness of analysis, while pathway RAG improves the precision of pathway retrieval. Their combined use allows scChat to achieve cutting-edge performance in contextualized single-cell analysis.

## Conclusions

In this study, we present scChat – a multi-agent scRNA-seq research co-scientist – that can autonomously generate executable plans for multi-step analyses, ranging from data preprocessing and follow-up analysis to results visualization. In this study, we present scChat – a multi-agent scRNA-seq research co-scientist – that reframes single-cell analysis as a conversational, plan-driven scientific process. scChat autonomously generates executable plans for multi-step analyses, spanning data preprocessing, downstream analysis, and results visualization. By coupling large language model–driven reasoning with established scRNA-seq analytical workflows, scChat introduces a generalizable co-pilot architecture that orchestrates cell annotation, differential expression, and functional interpretation in a unified and adaptive manner. Unlike traditional tools and function-specific models, scChat coordinates widely adopted packages such as Scanpy^3^, g:Profiler^41^, and GSEApy^42^ within an LLM-guided planning and execution framework, transforming conventional scRNA-seq pipelines into an interactive research process that supports context-aware, plain-language scientific inquiry. Beyond providing accurate annotation and enrichment, scChat can highlight biologically meaningful patterns that guide subsequent experimental design. To achieve this, we combined function calling with RAG, enabling the system to perform quantitative analysis while grounding its outputs in established databases. The capabilities of scChat are evaluated through cell type annotation accuracy test, failure mode testing, and ablation study. These tests highlight scChat’s robustness in delivering high cell type annotation accuracy and its ability to address scRNA-seq–specific questions in a fully automated, interactive manner. With excellent performance across diverse test datasets, scChat demonstrates its potential to serve as an AI co-scientist that accelerates and enriches biomedical research. However, key challenges remain: the incompleteness of biological knowledge bases can lead to missed cell type annotations, and the system’s reliance on user interaction for hypothesis generation highlights the limits of current multi-agent frameworks. Resolving these fundamental issues will be essential to unlocking the full potential of scChat and advancing the broader goal of AI-augmented scientific discovery. While scChat’s reliance on external LLM backends may introduce variability in conversational behavior over time, its underlying analytical computations remain deterministic and reproducible. Future work will focus on enabling fully local execution through fine-tuned open-source language models, thereby improving reproducibility, transparency, and data privacy. In parallel, we aim to explore more fine-grained evaluation protocols for open-ended biological reasoning, including approaches that separately characterize errors arising from knowledge-base incompleteness, dataset annotation mismatches, and multi-step reasoning failures. We view the development of principled and community-accepted evaluation frameworks for conversational biological analysis as an important open research direction beyond the scope of the present study.

## Methods

### [Multi-agent framework]

The multi-agent framework of scChat uses Langgraph^43^, a system for multi-agent design, as the backbone and comprises five agents: a planner, an executor, an evaluator, a critic, and a response generator. The entire workflow begins with the planner. Each agent’s function is described as follows: the **planner** searches for function execution and conversation history, parses the query, and decomposes it to generate a plan with several function calls arranged as steps in sequence. The **executor** performs the function specified in the plan iteratively. The **evaluator** validates the outcome of each function from the executor, handling errors and interrupting the plan to pass error messages to the response generator if needed. Additionally, it checks the availability of remaining steps and determines the next step in the workflow. The **critic** identifies potentially missing functions by creating a separate plan based on the function results, ensuring targeted analyses of specific cell types with all necessary downstream steps. Finally, the **response generator** compiles all relevant function results to generate the final response to the user’s query. After generating the response, it stores the final response and the function execution results in conversation and function histories, respectively. The hyperparameters for LLM can be figured out in Supplemental Information **Table S2**. The prompts that used for agent, on the other hand, are saved in **Fig. S14-18**.

### [scRNA-seq Dataset]

The scRNA-seq dataset is a primary input for scChat, which accepts data in .h5ad format, a common file type for storing single-cell RNA sequencing data. When multiple scRNA-seq samples are collected from different experiments, such as treatment and control groups, scChat requires users to upload an accompanying JSON file. This file matches the “sample id” label in the dataset’s observation layer and provides descriptions for each condition. These labels and descriptions help scChat distinguish changes between different conditions. This is especially useful when comparing gene expression, pathways, and cell populations, enabling scChat to offer valuable insights into the biological differences between groups.

### [Web Interface]

The web interface is the central hub for user interaction within the system, providing a seamless and intuitive platform for conducting complex scientific inquiries and analyses. Specifically, scChat performs all computational analysis on the user’s local machine and transmits only derived results and contextual summaries—rather than raw scRNA-seq data—to GPT-4o via the OpenAI API. The generated responses are then returned to the user through the interface. It allows users to input queries and upload various data types for comprehensive analysis. Integrated data visualization tools enable users to view graphs and charts directly within the interface, enhancing the interpretation of complex datasets. Additionally, export functionalities facilitate smooth transitions to external software, supporting further analysis and study.

### [Large Language Model (LLM)]

The LLM, GPT-4o, is crucial to the scChat platform, managing the execution of function calls, retrieval-augmented generation processes, and conversation history. To improve contextual analysis, all statistics generated by function calls, along with the research context, are stored in a vector database as part of the LLM’s context. This enables GPT-4o to access relevant information when answering follow-up questions, ensuring its analyses are based on actual data rather than speculation. The embedded statistics help anchor the LLM’s responses, reducing the risk of hallucination and increasing the reliability of the generated insights.

### [Retrieval-Augmented Generation (RAG) and context engineering]

Ensuring accuracy and relevance in LLMs, especially in specialized fields like scRNA-seq analysis, is difficult. Although LLMs generate text based on learned patterns, they can produce incorrect or superficial responses for complex tasks. RAG solves this by incorporating external, domain-specific knowledge, enabling the model to retrieve and reference relevant documents or databases.

In scChat, we use two methods to support RAG: the knowledge graph and vector database. The knowledge graphs, built with the Neo4j platform, store the cell type hierarchy system and an enrichment analysis database. During information retrieval from the knowledge graph, scChat uses structural Cypher prompts to communicate with the Neo4j database and gather the necessary information. The cell type hierarchy system consists of three nodes: “System,” “Organ,” and “CellType.” The “System” and “Organ” nodes are connected via “has organ” edges, and “System” links to related cell types through “has cell” edges. The “develops to” edge is used to connect the “Level” variable within each cell type, forming a cell lineage tree. In each “process cells” function, scChat uses system, organ, and cell type information to retrieve marker genes from the next-level cell type and conduct cell type annotation. The procedure that scChat retrieves information from the cell type knowledge graph, we call it cell type RAG. Notably, the cell type hierarchy system is built based on information rearranged from CellMarker^44^.

The pathway knowledge graph, on the other hand, contains three core nodes: “Database”, “GeneSetLibrary”, and “Pathway”. The “Database” encompasses all the enrichment methods that scChat can apply, including Gene Ontology (GO), Kyoto Encyclopedia of Genes and Genomes (KEGG), Reactome, and Gene Set Enrichment Analysis (GSEA). The first three databases are directly linked to their pathways with “found in” edges. After RAG retrieval, these three databases utilize g:Profiler^41^(version e113_eg59_p19_f6a03c19; corresponding to Ensembl release 113 and Ensembl Genomes release 59), a Python package, with cell type and database type as inputs to perform enrichment analysis. In contrast, GSEA needs to verify the desired gene set library with a “contains library” edge and then search for available pathways through a “contains pathway” edge. Subsequently, cell type and gene set library are used as inputs for GSEApy^42^ (v1.1.3) to perform enrichment analysis. To maximize coverage, scChat incorporates the complete collection of gene set libraries hosted by the GSEApy’s database, comprising 223 distinct libraries spanning transcription factor targets, drug perturbations, and cell-type signatures. All pathway and gene set terms derived from both g:Profiler–supported resources (GO, KEGG, and Reactome) and the full collection of GSEApy gene set libraries, together with their associated gene lists, are embedded into the vector database to enable semantic retrieval within the RAG framework.

Since the LLM relies heavily on the context window to remember past interactions, the conversation becomes more prone to hallucinations and incurs higher costs as more tokens are fed into the model over time. Storing all results in a vector database and retrieving only the necessary history can help address this problem. In scChat, we adopt a vector database that utilizes Hugging Face’s sentence transformers, specifically all-MiniLM-L6-v2^45^, as RAG to embed all the pathways, information from enrichment analysis, conversation history, and function execution history into chromaDB, an open-source embedding database, separately.

To facilitate semantic search, results from enrichment analysis, conversation history, and function history are embedded with specific metadata formats. For example, in enrichment analysis, the metadata includes term name, term ID, analyzed cell type, analysis type, sample condition, adjusted p-value, average log2 fold change, intersecting genes, intersecting size, rank, and term description. Conversation and function history use timestamps, session IDs, context, function names, and input parameters as metadata, respectively. This allows the planner and response generator to retrieve complete information by a simple keyword, reducing latency from semantic matching. Lastly, a pathway vector database is pre-built with the same setup, used to provide accurate pathway names and prevent semantic mismatches when retrieving information from the pathway knowledge graph. The entire process performed by scChat, including semantic search in the vector database and retrieval of function specifications from the knowledge graph, is referred to as pathway RAG.

In conclusion, the RAG system manages the knowledge graph and vector database, which not only improves scChat’s performance in cell type annotation and enrichment analysis but also provides scChat with a more effective way to remember previous interactions.

### [scRNA-seq data preprocessing and function calls]

LLMs such as GPT-4o, while powerful tools for generating insights and providing contextual understanding, are not designed to perform direct quantitative data analysis. This limitation arises because LLMs, although sophisticated, may hallucinate or produce outputs that are not strictly grounded in the underlying data. To address this challenge, scChat utilizes a series of predefined function calls, powered by the open-source package Scanpy^3^, to construct the necessary functions for analyzing raw scRNA-seq data. When users upload their raw scRNA-seq files to the platform and activate the chatbot, scChat automatically initiates a series of preprocessing steps followed by first-level cell type annotation. Specifically, scChat first performs standard cell- and gene-level quality control and computes common QC metrics, including the fraction of mitochondrially encoded transcripts, to identify and filter low-quality cells. Raw counts are then normalized to 10,000 per cell using cp10k normalization, followed by log1p transformation. Mitochondrial gene expression can contribute disproportionately to variance in a dataset-dependent manner, which may bias clustering toward stress-or quality-associated axes rather than lineage structure. In addition, mitochondrially encoded genes are frequently among the most highly expressed transcripts across diverse cell types, and their inclusion can cause mitochondrial signals to dominate short cluster-level gene summaries, thereby obscuring lineage-defining markers. For these reasons, scChat removes mitochondrially encoded genes, such as MT-prefixed genes, from the expression matrix prior to downstream feature selection and clustering. This task-driven design choice is intended to improve the specificity of large language model–based cell type annotation and is optimized for identity-centric atlas construction. We acknowledge that removing mitochondrial genes may diminish mitochondria-centered biological programs and influence variance structure as well as downstream feature selection and clustering in certain datasets. Future versions of scChat will consider providing user-facing options to flexibly include or exclude mitochondrial genes depending on analytical goals. Following preprocessing, scChat selects highly variable genes to enhance the discrimination of distinct cell types or states. Subsequently, a single-cell variational inference (scVI) model is trained on the processed scRNA-seq data under multiple conditions, which is used to mitigate batch effects and technical noise.

The “process cells” and “enrichment analysis” functions are two specialized functions that use the RAGs to retrieve external knowledge and perform specific tasks. The “process cells” function annotates cell types by combining external knowledge with expression profiles. First, based on the user query, it retrieves relevant child cell types and their marker genes from the cell type RAG. For example, within immune cells, child types include T cells, B cells, mononuclear phagocytes, and natural killer cells. Next, scChat cross-references this retrieved knowledge with the top 25 most highly expressed genes in each cluster. By matching cluster-specific gene expression with marker gene signatures, the function determines the most likely cell type label. The prompt used for cell type annotation is shown in **Fig. S11**. In scChat, enrichment analysis is conducted through a two-step retrieval process. First, instead of directly performing enrichment on the target cell type, scChat performs a semantic search within the pathway vector database to identify the top three pathways related to the query concept. These candidate pathways are then used as keywords to query the pathway knowledge graph. From the graph, scChat retrieves structured function specifications, such as the appropriate enrichment method and gene set library, which are then used as inputs to the enrichment function.

In contrast, other analytic functions, such as “dea split by condition”, which identifies differential gene expression with log2 fold change > 1 and adjusted p-value < 0.05, and “compare cell counts” that compares cell populations across conditions, do not require RAG participation. Similarly, visualization functions, including UMAP for cell type annotation, feature plots, violin and heat maps for gene expression, and bar or dot plots for enrichment analysis, are executed through the standard workflow. In these cases, the planner directly adds the required function to the plan, which then proceeds through the executor, evaluator, critic, and response generator without invoking the RAG. This highlights that RAGs are applied selectively during function execution, only when annotation or enrichment tasks demand external knowledge integration.

Each function is carefully designed to help researchers perform complex analyses and obtain meaningful insights from their data. By accepting various questions directly, the planner coordinates these functions, allowing researchers to concentrate on their scientific goals without needing to worry about specific function names or underlying code.

### [Intra-system interaction]

**Fig. 8** illustrates how scChat interacts with the four types of benchmarking questions, detailing the coordination among agents in answering them. **Fig. 8a** shows how scChat interacts with cell type identification questions. By providing the dataset’s metadata, such as the selected database (human or mouse), system, and organ, in the predefined instructional JSON file, and mentioning the target cell type in the user query, the cell type RAG returns candidate cell types and their marker genes for scChat to perform cell type annotation. Suppose the query involves identifying a cell type that is several levels away from the currently available cell types. In that case, the planner can recognize the issue and create a plan that quickly determines its existence by searching through various parent-child relationships. In the example, the query asks for CD4^+^ memory T cells. The planner starts by identifying the overlap between the parent cell type of the user’s target cell and the available cell types. In this case, the overlap is the immune cell. To reach the requested subtype, CD4^+^ effector memory T cell, the planner maps the lineage path from the immune cell down to the target. It then creates a stepwise plan, which the executor carries out. The evaluator iteratively verifies the presence of intermediate cell types, while the critic decides if additional refinement is needed. Finally, the response generator compiles the verified results into a user-facing reply and saves them for future use.

**Figure 8.**
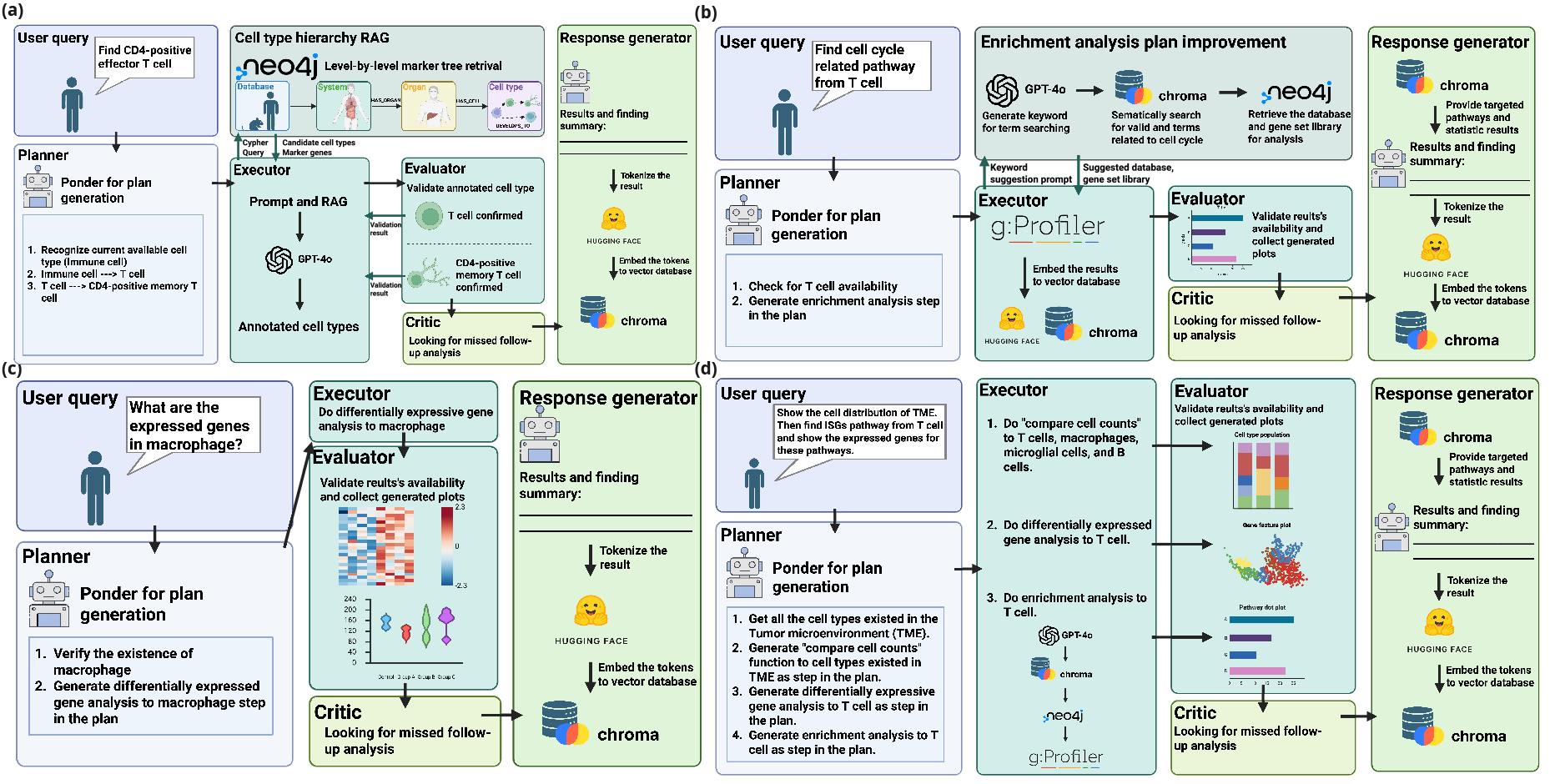
ScChat interactions with different types of questions (a) Cell type identification question. (b) Enrichment analysis question. (c) Differential gene expression question. (d) Complex query.

**Fig. 8b** explains how scChat interacts with enrichment analysis questions. It can be challenging to identify a target pathway without first selecting an analysis method, including GO, GSEA, KEGG, and Reactome, as well as the gene set library, which is only available when GSEA is chosen as the method. To address this, as the planner adds the enrichment analysis step to the plan, the pathway RAG enhances this step by specifying the method and gene set library containing the target term. For example, the user requests cell cycle-related terms from T cells. Based on this query, the planner then inserts the enrichment analysis step into the plan. Before the executor starts the analysis, the pathway RAG steps in to improve the plan by querying GPT-4o to retrieve the top three cell cycle-related pathways from the pre-built pathway vector database. These pathways serve as keywords, allowing the system to retrieve the corresponding method and gene set library from the knowledge graph. Once the plan is refined, the executor proceeds with the function and stores the enriched pathways in the results database. As the workflow reaches the response generator, instead of directly answering the question, it performs a semantic search in the results database. It consolidates all cell cycle-related pathways and their associated statistical values, including p-value, log2 fold change, and intersecting genes, to generate a response. This approach enables scChat to define function specifications precisely and to examine all enriched pathways, rather than being limited to the top enriched pathways.

**Fig. 8c** illustrates the situation in which differential gene expression analysis is necessary. To conduct a follow-up analysis, the planner first verifies if the target cell type exists, then adds a step to perform differential gene expression analysis on that cell type in the plan. In this case, differential gene expression in macrophages is used as an example. Next, the executor runs differential gene expression analysis on each sample condition of the macrophages, and the evaluator checks for errors. For differential gene expression, the user can request to generate a heat map, violin plot, or feature plot; the heat map is the default option. After executing the function, the workflow proceeds to the critic to assess the need for additional functions, followed by the response generator for response creation and storage.

A complex query in scChat involves function chaining rather than a single function call. As shown in **Fig. 8d**, the system first identifies that multiple steps are necessary, and the planner constructs a sequential plan accordingly. The executor then carries out each step, invoking the RAG for external knowledge retrieval when needed. At the same time, functions, such as differential gene expression and cell population comparison, can be executed directly. Finally, the evaluator, critic, and response generator review the outputs, decide if further refinement is required, and compile the final response.

### [Benchmarking pipeline and scoring metrics]

To systematically evaluate scChat across eight journal articles, specifically, a cancer-related dataset from Gondal et al^34^, a car T cell therapy dataset from Strati et al^35^, data on peripheral immune response for COVID-19 from Wilk et al^36^, immunophenotyping data of COVID-19 and influenza from Lee et al^37^, a retinal pigment epithelium dataset from Voigt et al^38^, a hematopoietic dataset from Triana et al^39^, a glioblastoma dataset from Bagley et al^26^, and a newborn human lung cell dataset from Bhattacharya^40^. We created an automated question–answer assessment pipeline. All the questions are designed to cover the scope of a predefined function or a typical workflow, from cell type identification to follow-up analyses, such as differential gene expression and enrichment analysis, as well as the ability to perform function chaining.

First, GPT-4o was prompted to produce question templates in four categories: cell type identification, differential gene expression, enrichment analysis, and complex query. Next, keywords were extracted from each article using GPT-4o and incorporated into the templates to develop article-specific benchmarking questions. To establish ground truth, the full journal articles were provided to GPT-4o, which generated reference answers for each question. Finally, GPT-4o evaluated scChat’s responses using a predefined rubric and scoring criteria, enabling consistent, large-scale assessment of chatbot performance. All the prompts, including the templates, prompt for question generation, prompt for evaluation, and prompt for agents are available in the supplementary information of **Fig. S10-13**.

To evaluate annotation accuracy, we use a hierarchical F1-score instead of exact label matching due to the complexity of cell lineage structures and variability in cell type naming across datasets. Model predictions are first mapped to the closest cell type in the CellxGene platform’s ontology, with the original dataset annotations serving as ground truth. Using these pairs, we calculate hierarchical recall and precision. To prevent inflation of precision from broader cell type prediction, we add a penalty factor (β) to the hierarchical F1 calculation. Full implementation details are available in the Supplemental Information **equation (1)-(8)**.

## Data availability

scChat is provided as an open-source software package at the GitHub repository https://github.com/li-group/scChat Several scRNA-seq samples and their research contexts are provided in the GitHub repository.

## Acknowledgements

We thank Bao and Li laboratory members for their technical assistance. This study was supported by startup funding from the Davidson School of Chemical Engineering and the College of Engineering at Purdue (X.B., C.L.), NIH NCI (grant no. R37CA265926 and R01CA293514 to X.B.), NSF (grant no. 2143064,2424004 and 2425704 to X.B. and C.L.), and Ralph W. and Grace M. Showalter Research Trust Grant (C.L.). This publication is also supported in part by Research Scholar Grant # RSG-25-1422119-01-IBCD from the American Cancer Society (X.B.). The authors also gratefully acknowledge support from the Purdue University Institute for Cancer Research, P30CA023168, Purdue Institute for Integrative Neuroscience (PIIN), Bindley Biosciences Center, and the Walther Cancer Foundation.

