## supplement for "scChat: A Large Language Model-Powered Co-Pilot for Contextualized Single-Cell RNA Sequencing Analysis"

#### Supplemental information

***Showcase study questions and responses***

**Table S1** Showcase study Q&A summary

| Question ID | Question | Concise Answer | Detailed Answer |
|---|---|---|---|
| Q01 | Which immune cell types dominate the TME before and after treatment? | Macrophages and T cells dominate; B cells are minor. | Based on the analysis results provided, the dominant immune cell types in the tumor microenvironment (TME) before and after treatment are as follows:<br><br>Pre-Treatment: T cells and macrophages dominate (notably abundant in patients 6 and 7). B cells are minor.<br>Post-Treatment: T cells remain dominant in P1, macrophages increase in P6, microglia increase in P7. B cells remain consistently low. |
| Q02 | What trends in T-cell abundance are observed across patients before and after treatment? | Overall unchanged; P1 increased; P6 and P7 decreased. | Patient 1: significant increase in T cells. Patient 6: decrease. Patient 7: decrease. Overall: variability across patients; some increase, some decrease. |
| Q03 | For each patient with paired samples, compare T-cell proportions before versus after treatment. | P1 increased; P6 and P7 decreased. | P1: increase in T cells; P6: decrease; P7: decrease. Reflects inter-patient differences in treatment response. |
| Q04 | How do myeloid populations (e.g., monocyte-derived macrophages and TREM2^hi microglia) shift after treatment across patients? | Only P1 shows concurrent monocyte-derived macrophages increase and TREM2^hi microglia decrease; others do not show this combined trend. | P1: macrophages increase, microglia decrease. P6: macrophages increase, microglia slightly increase. P7: macrophages decrease, microglia increase. Heterogeneous responses observed. |

| | | | |
|---|---|---|---|
| Q05 | Summarize the direction and magnitude of the myeloid shift in P1 (qualitative/approximate is sufficient). | monocyte-derived macrophages ~+15 percentage points; microglia ~−10 percentage points. | P1: macrophages increased, microglia decreased. Net shift suggests replacement of resident microglia by infiltrating macrophages. |
| Q06 | Are antigen-presentation programs upregulated in macrophages after treatment? Provide representative genes. | Yes; representative genes include HLA-DRA and CD74. | Yes. Antigen presentation genes (HLA-DRA, HLA-DRB1, HLA-DPB1, CD74) upregulated, indicating enhanced antigen presentation. |
| Q07 | Are interferon-stimulated gene signatures enriched in macrophages after treatment? | Yes, with moderate enrichment. | Yes. Interferon-stimulated genes moderately enriched, consistent with activated macrophage phenotype. |
| Q08 | Which major T-cell subpopulations are identified, and what roles might they play in the glioma TME before and after treatment? | Identifiable subpopulations include conventional/memory CD4 T, Tregs, resident-memory and effector/exhausted CD8 T, and IFN-stimulated T cells. | Tregs (FOXP3, IL2RA), central memory CD4 (IL7R, IL32), activated CD8 (CD69, PRKCQ), exhausted T cells (PDCD1 upregulation), IFN-stimulated T cells (IL32). Each plays immunosuppressive, supportive, cytotoxic, or inflammatory roles. |
| Q09 | Based on available expression data, did effector or exhaustion-related T-cell states change after treatment? | Yes; effector-exhausted T cells are more abundant post-treatment. | Effector-related genes (IL7R, NKG7, GZMK) upregulated post-treatment, indicating effector expansion. Exhaustion states less conclusive due to missing exhaustion marker data. |
| Q10 | Do T cells after treatment show enrichment of interferon-related gene expression programs (IFN-α/IFN-γ), and what does this imply for their functional state? | Yes, increased post-treatment, especially in CD8+ T cells; implies heightened inflammatory and activation. | Yes. IFN-γ response enriched, implying heightened inflammatory state. Suggests active anti-tumor response, but prolonged IFN exposure could lead to exhaustion. |

| | | | |
|---|---|---|---|
| Q11 | Among post-treatment which T subtype shows the strongest IFN-response program? | CD8+ T cells. | CD8+ effector memory T cells show strongest IFN-response (genes GZMK, NKG7, CCL5 upregulated). |
| Q12 | Which TCR-signaling or activation-related programs are upregulated in T cells after treatment, and what does this indicate about T-cell activation in the TME? | Evidence supports increased activation post-treatment (e.g., CD25, HLA-DR, CD38, MKI67 upregulated). | TCR signaling genes (CD3D, CD3E, ZAP70, ITK) and activation markers (CD25, CD69) upregulated. Indicates robust T-cell activation in TME post-treatment. |
| Q13 | Which CD4+ T-cell subtypes are present and which marker genes characterize them? | Mostly conventional effector/memory CD4 T (e.g., CD40LG, IL7R); Tregs are fewer (FOXP3, IL2RA/CD25). | Naive CD4 T cells (IL7R, TCF7, KLF2) and conventional alpha-beta CD4 T cells (IL32, CD40LG) dominate; Tregs fewer. |
| Q14 | In Tregs across patients, which immune function–related genes are highly expressed, and how might these genes contribute to immunosuppressive activity in the TME? | Tregs highly express FOXP3 and IL2RA (CD25). | FOXP3, IL2RA, CTLA4, TIGIT, BATF, GITR highly expressed. Together mediate immunosuppression by depleting IL-2, suppressing effector T-cell activation, and enhancing suppressive capacity. |
| Q15 | In CD8+ T cells across patients, which immune function–related genes are highly expressed, and what do they suggest about the cytotoxic or exhaustion states of these cells in the TME? | CD8+ T cells highly express GZMB, PRF1 (cytotoxicity) and PDCD1, LAG3 (exhaustion). | Cytotoxic genes (GZMA, GZMH, GZMK, NKG7, CCL5) and activation markers upregulated. Exhaustion marker evidence less clear, but cytotoxic profile is strong. |
| Q16 | What evidence suggests potential coordination between interferon-high macrophages and interferon-high T cells within the same TME? | A positive association is observed. | Both macrophages and T cells enriched for interferon-response pathways, with overlapping gene upregulation (IL32, GZMK). Suggests coordinated immune activation within TME. |

***LLM hyperparameter settings***

**Table S2** Hyperparameters for LLMs

| | Model | Temperature | Max token |
|---|---|---|---|
| Planner | GPT-4o (default) | 0.1 | Default |
| Pathway retrieving | GPT-4o-mini (default) | 0.1 | 50 |
| Cell type annotation | GPT-4o (default) | 0 | Default |
| Critic | GPT-4o (default) | 0 | Default |
| Critic question classifier | GPT-4o (default) | 0 | Default |
| Response generator | GPT-4o (default) | 0.3 | 800 |

***Cell type annotation scoring matrix***

$\forall v \in V,\ \mathrm{A}(v)$ is the set of ancestors of cell type v (1)

$\tau_i = \mathrm{A}(t_i)$, where $t_i \in V$ representing ground truth (2)

$P_i = \mathrm{A}(p_i)$, where $p_i \in V$ representing prediction (3)

$$\mathrm{h}P_i = \frac{|P_i \cap \tau_i|}{|P_i|}, \tag{4}$$

$$\mathrm{hP} = \frac{1}{N}\sum_{i=1}^{N} hP_i \tag{5}$$

$$\mathrm{h}R_i = \frac{|P_i \cap \tau_i|}{|\tau_i|}, \tag{6}$$

$$\mathrm{hR} = \frac{1}{N}\sum_{i=1}^{N} hR_i, \tag{7}$$

$$hF1 = (1+\beta^2)\cdot\frac{hP\cdot hR}{(\beta^2\cdot hP)+hR},\ \beta = 3 \tag{8}$$

***Cell type acronym list***

Astrocyte: AC

B cell: B cell

Activated CD8-positive, αβ T cell: $CD8^+$ αβ Tact

αβ T cell: αβ T cell

Basophil mast progenitor cell: BaMP

Blood vessel endothelial cell: bvEC

CD14-low, CD16-positive monocyte: $CD14^{low}$ $CD16^+$ Mo

CD14-positive monocyte: $CD14^+$ Mo

CD14-positive, CD16-positive monocyte: $CD14^+$ $CD16^+$ Mo

CD16-negative, CD56-bright NK: $CD16^{-}$ $CD56^{bright}$ NK
CD16-positive, CD56-dim NK: $CD16^{+}$ $CD56^{dim}$ NK
CD38-negative naive B cell: $CD38^{-}$ Bn
CD4-positive helper T cell: $CD4^{+}$ Th
CD4-positive naive T cell: $CD4^{+}$ Tn
CD4-positive effector T cell: $CD4^{+}$ Teff
CD4-positive, αβ cytotoxic T cell: $CD4^{+}$ αβ CTL
CD4-positive, αβ memory T cell: $CD4^{+}$ αβ Tm
CD4-positive, αβ T cell: $CD4^{+}$ αβ T
CD4-positive, αβ central memory T cell: $CD4^{+}$ αβ Tcm
CD4-positive, αβ effector memory T cell: $CD4^{+}$ αβ Tem
CD4-positive, αβ effector memory T cell, terminally differentiated: $CD4^{+}$ αβ Temra
CD8-positive effector T cell: $CD8^{+}$ Teff
CD8-positive effector memory T cell: $CD8^{+}$ Tem
CD8-positive, αβ cytotoxic T cell: $CD8^{+}$ αβ CTL
CD8-positive, αβ memory T cell: $CD8^{+}$ αβ Tm
CD8-positive, αβ memory T cell, CD45RO-positive: $CD8^{+}$ αβ Tm $CD45RO^{+}$
CD8-positive, αβ T cell: $CD8^{+}$ αβ T
CD8-positive, αβ thymocyte: $CD8^{+}$ αβ thymocyte
CD8-positive, αβ central memory T cell: $CD8^{+}$ αβ Tcm
CD8-positive, αβ effector memory T cell: $CD8^{+}$ αβ Tem
CD8-positive, αβ effector memory T cell, terminally differentiated: $CD8^{+}$ αβ Temra
Ciliated cell: CilC
Class-switched memory B cell: csBm
Classical monocyte: cMo
Common myeloid progenitor: CMP
Conventional dendritic cell: cDC
Conventional type 1 dendritic cell: cDC1
Conventional type 2 dendritic cell: cDC2
Cytotoxic T cell: CTL
Dendritic cell: DC
Early promyelocyte: Early ProMy
Endothelial cell: EC
Endothelial cell of pericentral hepatic sinusoid: EC (hepatic pericentral)
Enterocyte: EnteroC
Erythroblast: EryB
Erythrocyte: RBC
Erythroid lineage cell: EryLin

Erythroid progenitor cell: ErP
Exhausted T cell: Tex
Fibroblast: FB
Fibroblast of mammary gland: Mammary FB
Foveolar cell of stomach: FovC
GABAergic neuron: GABA neuron
γδ T cell: γδ T
Glial cell: GC
Glutamatergic neuron: Glu neuron
Goblet cell: GobC
Granulocyte-macrophage progenitor: GMP
Hematopoietic cell: HemC
Hematopoietic multipotent progenitor cell: MPP
Hematopoietic precursor cell: HPC
Hematopoietic stem cell: HSC
IgA plasma cell: IgA PC
IgG plasma cell: IgG PC
Immature B cell: ImmB
Inflammatory macrophage: Inflam MΦ
Innate lymphoid cell: ILC
Intermediate monocyte: iMo
Leukocyte: WBC
Lymphatic endothelial cell: LEC
Lymphocyte: Lymph
M1 macrophage: M1
M2 macrophage: M2
Macrophage: MΦ
Macrophage dendritic cell progenitor: MDP
Mast cell: MstC
Mature dendritic cell: mDC
Mature NK T cell: mNKT
Megakaryocyte: Mk
Megakaryocyte progenitor cell: MkP
Megakaryocyte-erythroid progenitor: MEP
Memory B cell: Bm
Memory T cell: Tm
Microglial cell: MG
Monocyte: Mo

Mononuclear phagocyte: MNP
Mucosal-associated invariant T cell: MAIT
Mural cell: MC
Myelocyte: MyeloC
Myeloid cell: Myeloid
Myeloid leukocyte: Myeloid WBC
Myofibroblast: MyoF
Naive B cell: Bn
Naive T cell: Tn
Naive thymus-derived CD4-positive, αβ T cell: $CD4^+$ αβ Tn (thymic)
Naive thymus-derived CD8-positive, αβ T cell: $CD8^+$ αβ Tn (thymic)
Natural killer cell: NK
Natural T-regulatory cell: nTreg
Neuron: Neuron
Neutrophil: Neu
Non-classical monocyte: ncMo
Oligodendrocyte: OLG
Paneth cell: PanC
Peptic cell: PepC
Pericyte: PeriC
Plasma cell: PlsmB
Plasmablast: PB
Plasmacytoid dendritic cell: pDC
Platelet: PLT
Pro-B cell: Pro-B
Progenitor cell: ProgC
Promonocyte: ProMo
Promyelocyte: ProMy
Radial glial cell: RGC
Regulatory T cell: Treg
Smooth muscle cell: SMC
Stem cell: StemC
Stromal cell: SC
T-helper 1 cell: Th1
T-helper 2 cell: Th2
T-helper 17 cell: Th17
Transitional stage B cell: TransB
Unswitched memory B cell: usBm

Vascular endothelial cell: vEc
Arterial endothelial cell: aEC
Capillary endothelial cell: cEC
Venous endothelial cell: vECn
Alveolar macrophage: AM
Kupffer cell: KC
Hofbauer cell: HBC
Kidney-resident macrophage: kMΦ
Intestinal macrophage: iMΦ
Cycling cell: Cycling
Mitochondrial-associated cell: Mito
Ribosomal-associated cell: Ribo
Heat shock–associated cell: HSP
Common lymphoid progenitor: CLP
Lymphoid-primed multipotent progenitor: LMPP
Early T lymphoid progenitor: ETP
Metanephric mesenchyme stem cell: MMMSC
Neuroepithelial cell: NEC
Tuft cell (intestinal): Tuft
BEST4-positive enterocyte: $BEST4^{+}$ Ent
Enteroendocrine cell: EEC
Intestinal stem cell: ISC
M cell (microfold cell): MC
Podocyte: Podo
Mesangial cell: MesC
Parietal epithelial cell: PEC
Juxtaglomerular cell: JG
Macula densa epithelial cell: MD
Glomerular capillary endothelial cell: gEC
Proximal tubule epithelial cell: PT
Proximal convoluted tubule epithelial cell: PCT
Distal convoluted tubule epithelial cell: DCT
Thin descending limb epithelial cell (Loop of Henle): tDL
Thin ascending limb epithelial cell (Loop of Henle): tAL
Thick ascending limb epithelial cell (Loop of Henle): TAL
Collecting duct epithelial cell: CD
Collecting duct principal cell: CD-PC
Collecting duct intercalated cell: CD-IC

Collecting duct alpha-intercalated cell: CD-αIC

Collecting duct beta-intercalated cell: CD-βIC

Connecting tubule epithelial cell: CNT

Nephron epithelial cell: NephEpi

Nephron progenitor cell: NPC

Peritubular capillary endothelial cell: ptEC

##### *T cell and macrophage supplementary*

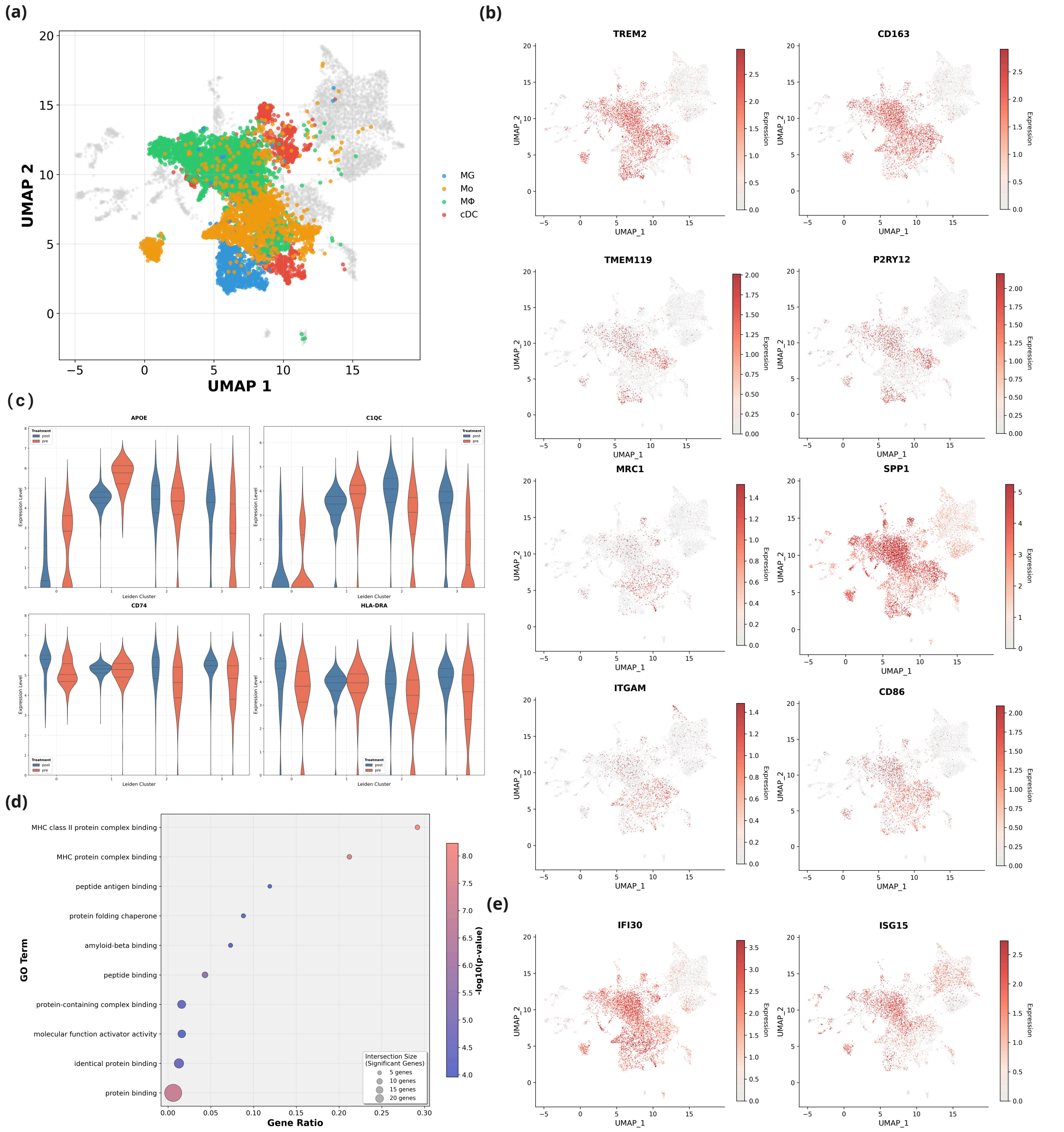


**Figure S1.** Bioinformatic analysis of macrophage-like cell subsets reveals increased myeloid inflammation and enhanced antigen-presenting phenotype. (a) UMAP visualization of macrophage-like cells from the tumor microenvironment (TME) of three paired patients (Pre and Post CAR T-cell infusion; six samples in total), after subsetting myeloid cells from the global UMAP shown in **Fig. 2b**. (b) Expression of selected marker genes for each cluster, displayed as feature plots, as in **Fig. 2b**. (c) Violin plots showing expression of macrophage state–associated genes across clusters in Pre versus Post samples, alongside the relative abundance of each cluster within each patient. (d) Gene ontology (GO) enrichment analysis of macrophage-like cells defined in (a).

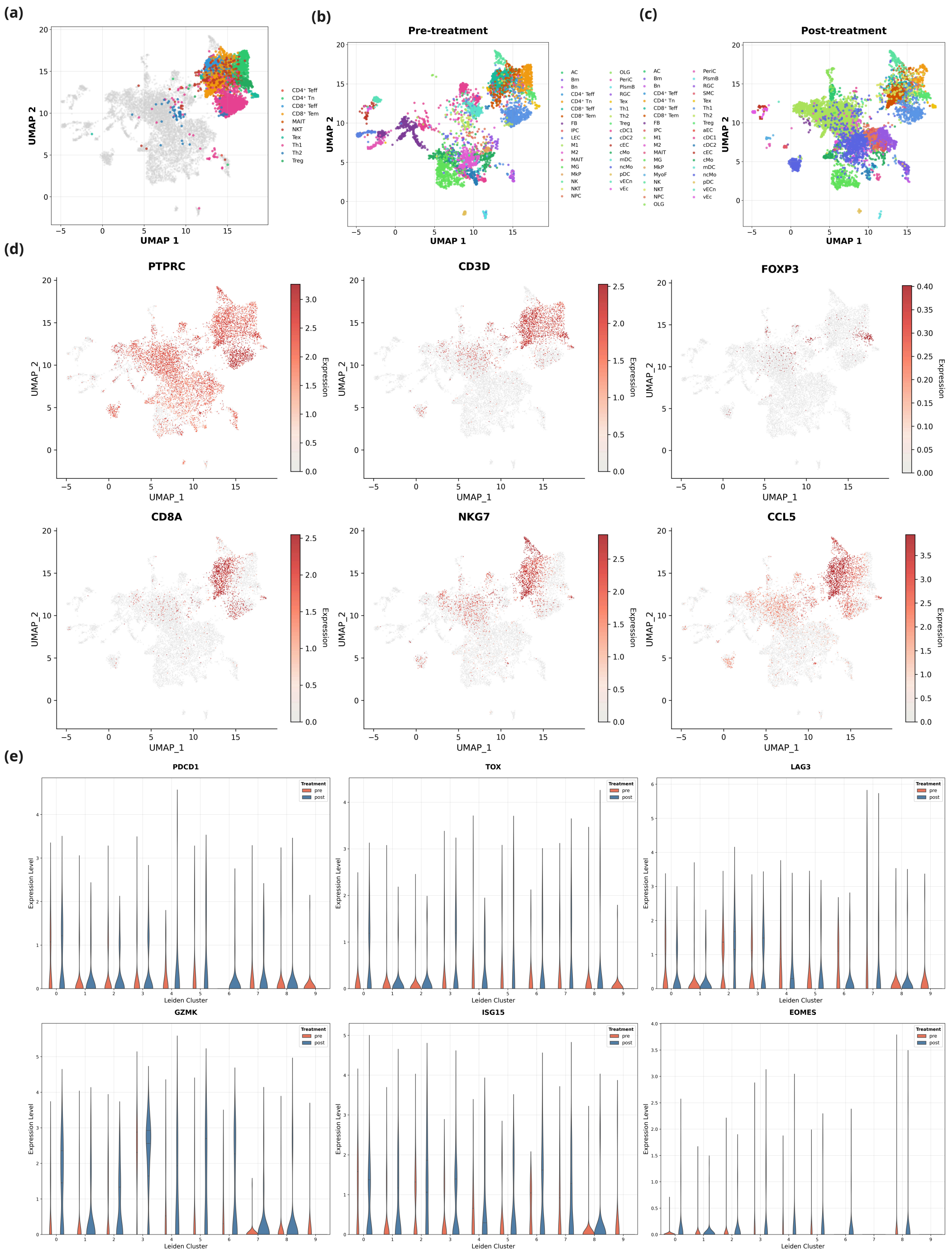


**Figure S2.** Bioinformatic analysis of T-cell subsets reveals increased T-cell exhaustion and interferon-stimulated T cells after treatment in pancreatic cancer. (a) UMAP visualization of T cells from the tumor microenvironment (TME) of three paired patients (Pre and Post CAR T-cell infusion; six samples in total), after subsetting from the global UMAP shown in **Fig. 2b**. (b) and (c), UMAP visualization of the overall TME cell populations from the same three paired patients, displayed separately for Pre (left) and Post (right) treatment. (d),Expression of selected T cell–related marker genes across clusters, shown as feature plots as in **Fig. 2b.** (e) Violin plots showing expression of T cell exhaustion– and IFN-response–associated genes across clusters in Pre versus Post samples, alongside the relative abundance of each cluster within each patient.

### *Cell type annotation comparisons*

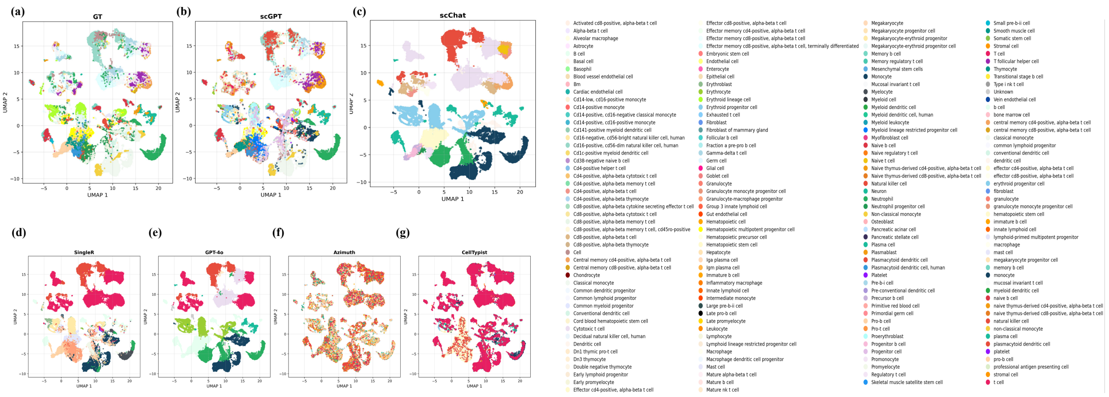


**Figure S3.** Cell type annotation for the bone marrow dataset

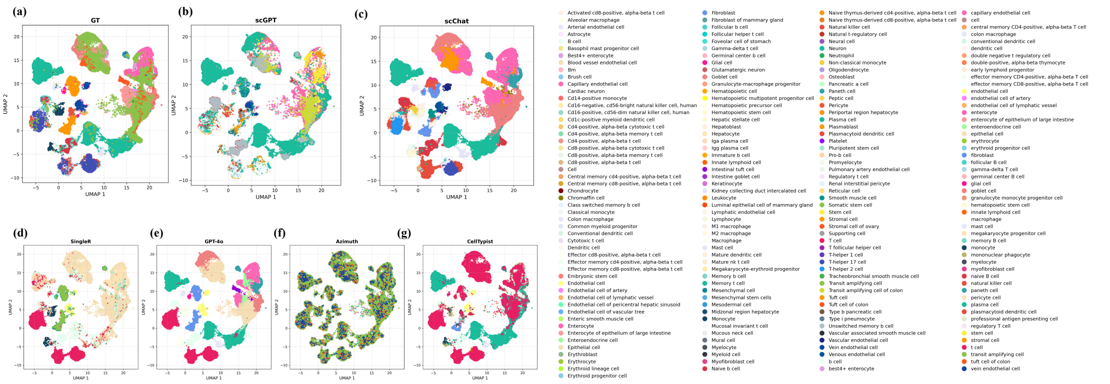


**Figure S4.** Cell type annotation for the intestine dataset

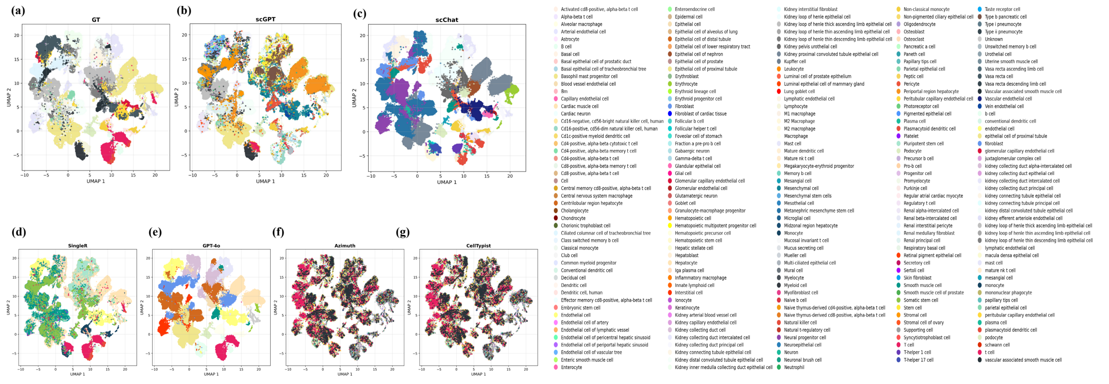

**Figure S5.** Cell type annotation for the kidney dataset

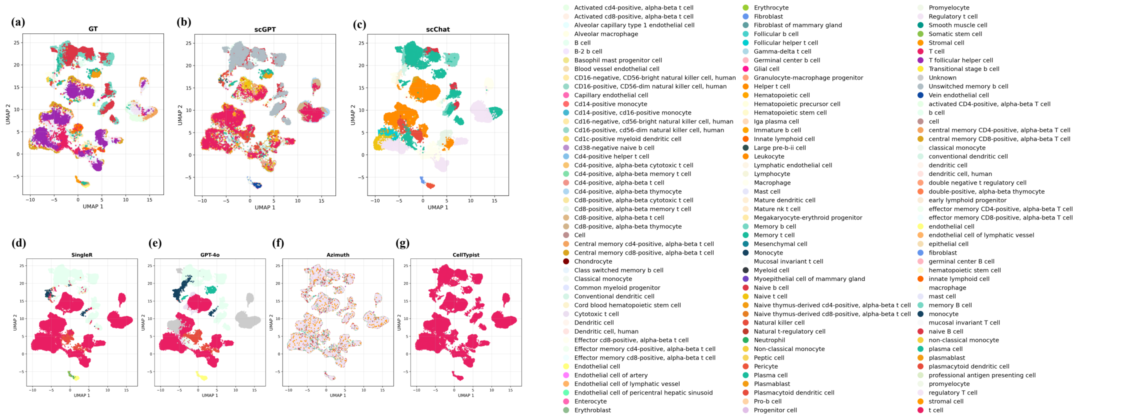


**Figure S6.** Cell type annotation for the lymph node dataset

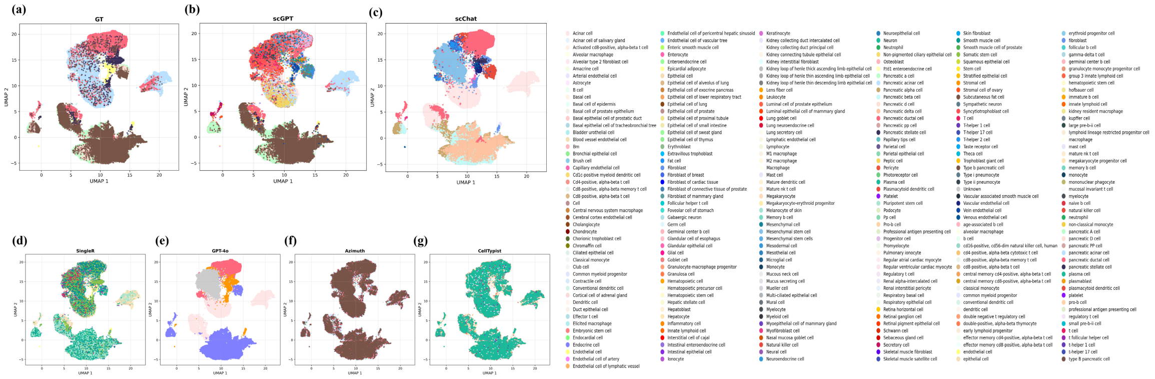


**Figure S7.** Cell type annotation for the pancreas dataset

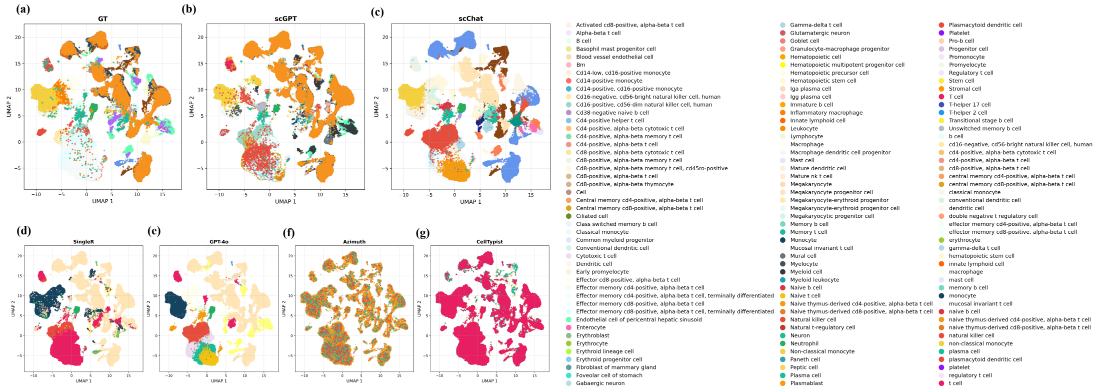


**Figure S8.** Cell type annotation for the liver dataset

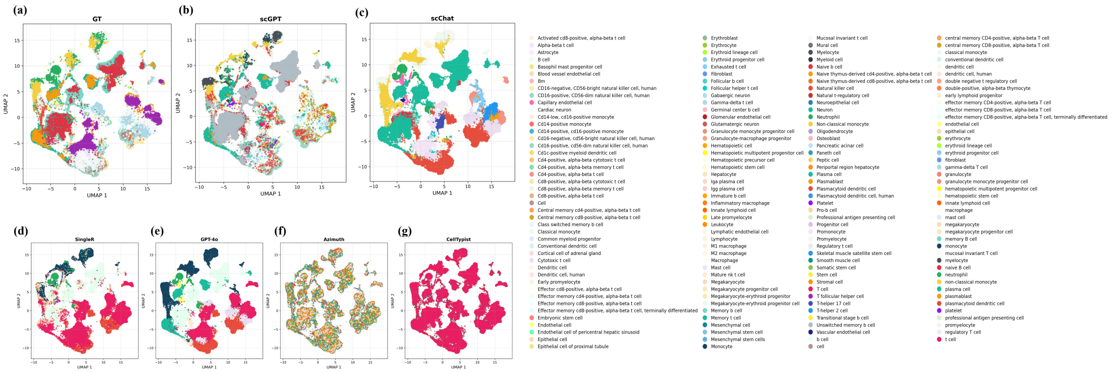


**Figure S9.** Cell type annotation for the spleen dataset

#### *Prompt engineering for cell annotation and benchmarking*

##### Cell type identification

1. How can we differentiate between {cell_type_A} and {cell_type_B} using their canonical markers?
2. How is the known cell type '{cell_type_A}' distinguished from the '{cell_type_B}' population, which is characterized by markers like {marker}?

##### Differentially expressed gene

1. Regarding the {cell_type_plural}, what {change_type} in {gene_name} expression was observed in {specific_context} when comparing {condition1} to {condition2}?
2. Regarding the {cell_type_plural}, what changes in {gene_name} expression were observed in {specific_context} when comparing {condition1} to {condition2}?

##### Enrichment analysis

1. Can we confirm if the {pathway_name} is enriched in {cell_type} from the {patient_group}?
2. The pathway analysis for the known '{cell_type}' population from the {context} displays what kind of key observation regarding the '{signature_name}' signature, including its significance?

##### Complex query

1. What is the relationship between the abundance of {cell_type} and the activity of {pathway/process}, and how might this impact clinical outcomes in this study's findings?
2. Let's synthesize the findings to explain how the interplay between {cell_types_list} and {pathway/process} contributes to the overall {environment/outcome}.
3. What are the key functional implications of {element} abundance changes observed in the experimental conditions?
4. Based on the successful identification of {cell_type}, the enrichment of the {pathway_element}, and the expression changes in {genes}, synthesize a comprehensive mechanistic explanation for how these elements interact to drive the observed therapeutic response across {conditions}.
5. Integrate the findings from {cell_type} populations, {pathway} activity, and {expression_patterns} to predict the long-term clinical implications and identify the most promising therapeutic targets for future intervention strategies.
6. Based on the {cell_type} population analysis and {pathway} activity, what is the primary mechanistic relationship that explains the therapeutic response observed in this study's findings?
7. How does the interaction between {cell_type} and {pathway_element} contribute to the overall treatment outcome, and what are the key regulatory factors involved?
8. What is the primary observation regarding {cell_type} behavior in the experimental conditions?
9. Describe the main finding related to {cell_type} in the experimental conditions and its implications for treatment efficacy.

**Figure S10.** Benchmarking question templates

**Cell type annotation prompt**

You are a bioinformatics researcher that can do the cell annotation.
The following are the data and its decription that you will receive.
* 1. Gene list: top 25 cells arragned from the highest to lowest expression levels in each cluster.
* 2. Marker tree: marker genes that can help annotating cell type.

Identify the cell type for each cluster using the following markers in the marker tree.
This means you will have to use the markers to annotate the cell type in the gene list.
Provide your result as the most specific cell type that is possible to be determined.

Provide your output in the following example format:
Analysis: group_to_cell_type = {'0': 'Cell type','1': 'Cell type','2': 'Cell type','3': 'Cell type','4': 'Cell type', ...}.

Strictly adhere to following rules:
* 1. Adhere to the dictionary format and do not include any additional words or explanations.
* 2. The cluster number in the result dictionary must be arranged with raising power.
* 3. The annotated cell type must exist in the marker tree.

**Figure S11.** Cell type annotation prompt

**Evaluation prompt**

You are an expert AI-powered evaluation model designed to evaluate systems that should analyze provided datasets.
Your evaluation must HEAVILY FAVOR answers that demonstrate actual data analysis over pre-trained knowledge.
**CORE PRINCIPLE**: This evaluation is for data analysis systems. An answer with actual statistical results from data analysis should ALWAYS score higher than textbook knowledge.
**CRITICAL EVALUATION FRAMEWORK:**
**Data Analysis Evidence Detection (PRIORITY #1)**:
**STRONG ANALYSIS EVIDENCE** (+2 bonus points):
- Specific statistical values (log fold change, p-values, adjusted p-values)
- References to analysis functions or methods
- Sample-specific results (P1, P6, P7 data)
- Differential expression analysis results
- Dataset-derived conclusions with quantitative support

**MODERATE ANALYSIS EVIDENCE** (+1 bonus point):
- References to single-cell RNA-seq analysis
- Mentions of data processing or analysis steps
- Context-specific findings that suggest data usage

**NO ANALYSIS EVIDENCE** (-2 penalty points):
- Pure textbook/general knowledge without data specificity
- Statements like "typically," "generally," "based on literature"
- Admissions of inability to access data
- Generic biological explanations without dataset grounding

**SCORING RUBRIC (0-5):**
**5 (Excellent)**: Strong analysis evidence + accurate findings. May differ from ground truth if analysis is well-supported.
**4 (Good)**: Clear analysis evidence + mostly accurate. Minor deviations acceptable if data-supported.
**3 (Acceptable)**: Some analysis evidence OR accurate ground truth coverage without analysis.
**2 (Poor)**: Minimal analysis evidence + significant ground truth deviations, OR pure knowledge-based answer when analysis was required.
**1 (Very Poor)**: No analysis evidence when required + inaccurate information.
**0 (No Value)**: Explicitly refuses to analyze data OR completely wrong when analysis was clearly required.

**EVALUATION INSTRUCTIONS:**
1. **First, identify if the question requires data analysis** by looking for:
- Expression level queries
- Comparison across conditions/patients
- Pathway analysis requests
- Population-specific findings
2. **Assess analysis evidence level** using the framework above
3. **Apply scoring bonuses/penalties** based on analysis evidence
4. **Compare to ground truth** while considering:
- If strong analysis evidence exists, accept valid scientific findings but add penalty if different from ground truth
- If no analysis evidence, require strict ground truth adherence

**Figure S12.** Automatic evaluation prompt

**Question generation prompt**

You are a biomedical research analyst specializing in extracting detailed molecular and cellular information from scientific papers AND generating evaluation questions based on templates.

#### INFORMATION EXTRACTION
CRITICAL: Extract EVERY gene mentioned, including:
- Markers (CD34, CD19, CD3, CD4, CD8, CD56, CD14, CD16, CD33, CD38, CD45, CD117, CD123, CD326, CD71, etc.)
- Transcription factors (GATA1, PU.1, RUNX1, CEBPα, PAX5, etc.)
- Differentiation markers (Tim3, CD11a, CD11c, CD98, etc.)
- Functional proteins (BCL2, MYC, FLT3, NPM1, etc.)
- Cytokines and receptors (IL2, IL7, CXCR4, etc.)
- ANY gene symbol in ANY context

For gene_details, ALWAYS provide condition_comparison when possible:
- If the paper mentions "expressed in X cells" → ["negative", "positive"]
- If mentioned as "marker for Y" → ["non-Y cells", "Y cells"]
- If "identified by FISH/IHC" → ["unstained", "stained"]
- If "upregulated/increased" → ["baseline", "activated"]
- If "downregulated/decreased" → ["baseline", "suppressed"]
- If comparing tissues → ["normal tissue", "diseased tissue"]
- If comparing time points → ["early", "late"]

Return extraction as JSON:
```
{
    "cell_types_identified": [...],
    "gene_details": [...],
    "gene_sets_pathways": [...],
    "key_observations": [...]
}
```
#### QUESTION GENERATION
##### Question Generation Rules:
- Generate 3-5 questions per template category where sufficient data exists
- Fill template placeholders with actual entities from the paper
- Ensure questions are answerable based on the paper content
- Include specific context (patient groups, conditions, markers)
- Create ground truth answers based on extracted information

##### Template Categories:
- **cell_type_identification**: Questions about distinguishing cell types using markers
- **differentially_expressed_gene**: Questions about gene expression changes between conditions
- **enrichment_analysis**: Questions about pathway/gene set enrichment
- **complex_query**: Multi-part questions requiring synthesis of findings

##### Output Format for Questions:
```
{
    "questions": [
        {
            "question_id": "Q1",
            "category": "cell_type_identification",
            "question": "How can we differentiate between B cells and T cells using their canonical markers?",
            "ground_truth": "B cells are identified by CD19, CD20, and CD79a markers, while T cells are characterized by CD3, CD4, and CD8 markers."
            }
        }
    ]
}
```
IMPORTANT:
- ALWAYS include condition_comparison for gene_details
- Ensure generated questions have factual, paper-based ground truth answers
- Include empty arrays [] only if truly no data exists
- Questions should test understanding of the paper's specific findings, not general knowledge

**Figure S13.** Benchmarking question generation prompt

**Planner prompt**

```
You are an intelligent planner for single-cell RNA-seq analysis.
Create a step-by-step execution plan (JSON) for the user query. CONTEXT:
- Available cell types ({len(available_cell_types)}): {', '.join(sorted(available_cell_types))}
- Failed discoveries: {', '.join(sorted(unavailable_cell_types)) if unavailable_cell_types else "None"}
- Status: {'Expanded' if len(available_cell_types) > len(self.initial_cell_types) else 'Initial only'}
- Previous analyses: {json.dumps(function_history, indent=2)} {"- Conversation context: " + conversation_context if conversation_context else ""}
Available functions: {self._summarize_functions(available_functions)} User question: "{message}"
DECISION LOGIC:
1. GENE/MARKER (differences, markers, counts) → DEA (dea_split_by_condition, compare_cell_counts)
2. PATHWAY (functional, ontology, signatures) → Enrichment (perform_enrichment_analyses)
3. CELL ABUNDANCE (proportions, quantities) → Counting (process_cells, compare_cell_counts, stacked plots)
4. VISUALIZATION ONLY (e.g. "show plot") → Set "visualization_only": true.
5. CONVERSATIONAL → No analysis steps.
JSON FORMAT:
{{
    "plan_summary": "Brief strategy description",
    "visualization_only": true/false,
    "steps": [
        {{ "step_type": "analysis|visualization|conversation",
           "function_name": "exact_name",
           "parameters": {{"param": "value"}},
           "description": "Step goal",
           "expected_outcome": "Step result",
           "target_cell_type": "Cell type name"
        }}
    ]
}}
CRITICAL RULES:
1. CELL TYPES: Only plan for "Available cell types". If missing, discover first (process_cells). Skip unavailable types.
2. GRANULARITY: Analyze one cell type per step (e.g., Step 1: T cell, Step 2: B cell). Never use list "T cells, B cells".
3. ORDER: "dea_split_by_condition" MUST precede "display_dea_heatmap" for the same cell type.
TOOL INSTRUCTIONS: - Enrichment (perform_enrichment_analyses):
* MINIMAL params: {{"cell_type": "Name"}}. NO "analyses" or "pathway_include" (auto-detected).
-  Visualizations: * Enrichment: "display_enrichment_visualization" (analysis="go"|"gsea", plot_type="bar"|"dot"|"both").
* Cell Counts (Aggregate): "display_cell_count_comparison" (after multiple compare_cell_counts).
Cell Counts (Stacked): "display_cell_count_stacked_plot" (for clinical/treatment groups, list all cell_types).
* DEA Heatmap: "display_dea_heatmap" (IMMEDIATELY after DEA step).
* UMAP (Gene): "display_feature_plot" (genes=["A","B"]).
* UMAP (Cluster): "display_leiden_umap" (shows cluster structure).
* UMAP (Global): "display_overall_umap" (color_mode="cell_type"|"accumulative_leiden").
* Expression (Dist): "display_violin_plot" (compare conditions, params: cell_type, genes).
- Semantic Search (search_enrichment_semantic):
* Use for specific pathway queries not in top results or context-based search. * Params: query="term", cell_type="Name".
```

**Figure S14.** Planner prompt

**Critic question classifier prompt**

```
User asked: "{original_question}"
The analysis was planned for '{unavailable_cell_type}' but this cell type was not found in the dataset.
Available cell types: {', '.join(sorted(available_cell_types))}
Should we analyze any of the available cell types instead to answer the user's question?
Consider biological relevance and semantic similarity.
If yes, return the most appropriate cell type name exactly as it appears in the available list.
If no suitable alternative, return "none".
Cell type or "none":
```

**Figure S15.** Critic question classifier prompt

##### Response generator prompt

**System Message:**
You are a single-cell RNA-seq analysis expert who provides direct, specific answers to user questions based on analysis results.

**User Message Template:**
You are a single-cell RNA-seq analysis expert.
USER'S QUESTION: "{original_question}"
[If Question Type & Guidance Provided]
QUESTION TYPE: {question_type}
RESPONSE GUIDANCE: {guidance}
[If Relevance Categories Provided]
ANALYSIS RELEVANCE:
- Primary analyses (focus on these): {relevance_categories.get("primary", [])}
- Secondary analyses (use for support): {relevance_categories.get("secondary", [])}
[If Conversation Context Provided]
CURRENT SESSION CONTEXT:
{conversation_context}
Please use the current session information above to provide accurate, context-aware responses.
CURRENT ANALYSIS RESULTS:
{formatted_findings}
[If Failed Analyses]
FAILED ANALYSES:
{failed_analyses_list}
INSTRUCTIONS:
1. Answer the user's question directly using the available data
2. Use specific gene names, pathways, and statistics from the results
3. If analyses failed, acknowledge this but provide insights using available data and biological knowledge
4. For comparisons, list concrete distinguishing features with specific molecular evidence
5. Be comprehensive but concise
6. Focus on answering the specific question asked, not providing general information
[If Conversation Context Provided]
7. Consider the conversation history and maintain continuity
8. Reference specific previous discussions when relevant
9. If the user is referring to something from earlier, address it specifically
[If Discovery Context Provided]
CELL TYPE DISCOVERY INFORMATION:
{discovery_context}
CRITICAL: If the user asks to "find" a cell type and you see it was discovered above,
lead with " [Cell Type] successfully discovered!" and explain it's now available for analysis.

Answer:

**Figure S16.** Response generator prompt

##### Pathway RAG retrieving prompt

Analyze this enrichment analysis query and determine intent:
User Query: "{message}"
First classify the user intent:
1. EXPLICIT_METHOD: User explicitly requests specific analysis methods
Examples: "run GO analysis", "perform KEGG", "do GSEA with MSigDB_Hallmark_2020"
2. PATHWAY_SPECIFIC: User mentions specific biological pathways/processes
Examples: "find interferon response pathways", "cell cycle regulation", "apoptosis pathways"
3. GENERAL: General enrichment without specific methods or pathways
Examples: "do enrichment analysis", "find enriched pathways"
Response format:
- For EXPLICIT_METHOD: Return "EXPLICIT_METHOD:method1,method2" or "EXPLICIT_METHOD:gsea:LibraryName"
- For PATHWAY_SPECIFIC: Return only the pathway keywords (like before)
- For GENERAL: Return empty string
Examples:
- "run GO analysis" → "EXPLICIT_METHOD:go"
- "perform KEGG and Reactome" → "EXPLICIT_METHOD:kegg,reactome"
- "GSEA with MSigDB_Hallmark_2020" → "EXPLICIT_METHOD:gsea:MSigDB_Hallmark_2020"
- "find interferon response pathways" → "interferon response"
- "do enrichment analysis" → ""
Extract specific pathway terms, biological processes, or cellular functions mentioned.
Return only the most relevant pathway keywords as a single string.
If no specific pathways are mentioned, return an empty string.
Result:

**Figure S17.** Pathway RAG retrieving prompt

#### Critic prompt

You are analyzing what bioinformatics analyses are needed for a specific cell type.
User Question: "{original_question}"
Cell Type: "{cell_type}"
Question Type: {question_type}
Available analysis functions with detailed descriptions:
CORE ANALYSIS FUNCTIONS:
- perform_enrichment_analyses: Run enrichment analyses on DE genes for a cell type. Supports REACTOME (pathways), GO (gene ontology), KEGG (pathways), GSEA (gene set enrichment).
- Use for pathway analysis when user asks about biological processes, pathways, or gene function.
- dea_split_by_condition: Perform differential expression analysis (DEA) split by condition. Use when comparing conditions or when user asks about gene expression differences between experimental groups.
- compare_cell_counts: Compare cell counts between experimental conditions for specific cell types. Use when analyzing how cell type abundance differs across conditions (e.g., pre vs post treatment, healthy vs disease).
VISUALIZATION FUNCTIONS:
- display_enrichment_visualization: PREFERRED function for showing comprehensive enrichment visualization with both barplot and dotplot. Use after running enrichment analyses to visualize results.
- display_feature_plot: Display gene expression on UMAP for specific genes. Use when user asks to visualize specific gene expression patterns or wants to see where certain genes are expressed.
- display_violin_plot: Display violin plots showing gene expression across leiden clusters with treatment comparison. Use when user wants to compare gene expression distributions across clusters or conditions.
- display_processed_umap: Display UMAP with cell type annotations. Use when user wants to see cell type annotations on UMAP.
- display_leiden_umap: Display UMAP colored by leiden clustering. Use when user wants to see clustering results or cluster-based analysis.
- display_overall_umap: Display overall UMAP with various coloring options (cell_type, leiden, sample). Use for general UMAP visualization requests.
- display_dotplot: Display dotplot for annotated results. Use when user wants to see gene expression patterns across cell types.
- display_dea_heatmap: Display DEA heatmap showing top differentially expressed genes. Use when user wants to see differential expression results as a heatmap.
- display_cell_count_stacked_plot: Display stacked bar plots of cell type counts across conditions. Use when user wants to see cell type abundance or proportions across experimental conditions.
SEARCH FUNCTIONS:
- search_enrichment_semantic: Search all enrichment terms semantically to find specific pathways or biological processes.
- Use when user asks about specific pathways, terms, or biological processes that might not appear in standard top results.
Task: Determine which analyses are needed for {cell_type} to answer the user's question.
IMPORTANT: The cell type "{cell_type}" already exists in the dataset. DO NOT suggest process_cells for this cell type.
Consider based on question type:
1. Gene/marker questions (differentiate X from Y, markers of X, gene expression) → Use dea_split_by_condition + display_dea_heatmap
2. Pathway/biological process questions → Use perform_enrichment_analyses + search_enrichment_semantic + display_enrichment_visualization
3. Cell abundance questions → Use compare_cell_counts + display_cell_count_stacked_plot
4. Specific gene expression questions → Use display_feature_plot or display_violin_plot
5. UMAP/clustering visualization requests → Use display_processed_umap, display_leiden_umap, or display_overall_umap
6. Specific pathway search → Use search_enrichment_semantic
7. If the question is about finding cell types (requires process_cells) → Return empty array []
CRITICAL GUIDELINES FOR GENE/MARKER QUESTIONS:
- For gene/marker questions, use dea_split_by_condition + display_dea_heatmap
- DO NOT include enrichment analyses for marker identification
- DO NOT suggest process_cells for already-available cell types
- Include relevant visualization functions for gene expression results
CRITICAL GUIDELINES FOR CELL TYPE DISCOVERY QUESTIONS:
- If the question is about finding/discovering cell types that require process_cells, return empty array []
- The critic should NOT suggest analyses for cell types that don't exist yet
- Cell type discovery is handled by the planner, not the critic
VISUALIZATION GUIDELINES:
- For enrichment results: display_enrichment_visualization
- For gene expression: display_feature_plot, display_violin_plot, or display_dea_heatmap
- For cell type visualization: display_processed_umap, display_leiden_umap, display_overall_umap
- For cell counts: display_cell_count_stacked_plot
- For gene patterns across cell types: display_dotplot
GENERAL GUIDELINES:
- Include appropriate visualization functions based on the analysis type
- Only include display_enrichment_visualization ONCE per cell type
- Choose the most relevant UMAP function based on what user wants to see
- Return ONLY a valid JSON array of function names, nothing else
Example responses:
- Pathway question: ["perform_enrichment_analyses", "search_enrichment_semantic", "display_enrichment_visualization"]
- Marker gene question: ["dea_split_by_condition", "display_dea_heatmap"]
- Gene expression visualization: ["display_feature_plot"]
- Cell type UMAP: ["display_processed_umap"]
- Cell abundance: ["compare_cell_counts", "display_cell_count_stacked_plot"]
Required analyses for {cell_type}:

**Figure S18.** Critic prompt
